# Mapping human natural killer cell development in pediatric tonsil by imaging mass cytometry and high-resolution microscopy

**DOI:** 10.1101/2023.09.05.556371

**Authors:** Everardo Hegewisch-Solloa, Janine E. Melsen, Hiranmayi Ravichandran, André F. Rendeiro, Aharon G. Freud, Bethany Mundy-Bosse, Johannes C. Melms, Shira E. Eisman, Benjamin Izar, Eli Grunstein, Thomas J. Connors, Olivier Elemento, Amir Horowitz, Emily M. Mace

## Abstract

Natural killer (NK) cells develop from CD34+ progenitors in a stage-specific manner defined by changes in cell surface receptor expression and function. Secondary lymphoid tissues, including tonsil, are sites of human NK cell development. Here we present new insights into human NK cell development in pediatric tonsil using cyclic immunofluorescence and imaging mass cytometry. We show that NK cell subset localization and interactions are dependent on NK cell developmental stage and tissue residency. NK cell progenitors are found in the interfollicular domain in proximity to cytokine-expressing stromal cells that promote proliferation and maturation. Mature NK cells are primarily found in the T-cell rich parafollicular domain engaging in cell-cell interactions that differ depending on their stage and tissue residency. The presence of local inflammation results in changes in NK cell interactions, abundance, and localization. This study provides the first comprehensive atlas of human NK cell development in secondary lymphoid tissue.

## Introduction

Human natural killer (NK) cell development can be defined by 6 stages characterized by their lineage potential, cell surface protein expression, and function (1–3). These stages can be described by their cell surface receptor expression and lack of T cell (CD3), B cell (CD19), and myeloid cell (CD14) lineage markers. Stage 1 is a multipotent lymphoid progenitor defined as being CD34+CD45RA+CD49d+integrin β7+CD117–; stage 2 is CD34+CD117+; stage 3 is an innate lymphocyte precursor defined as being CD117+CD127+CD94–; stage 4 is an NK committed progenitor that is CD56^bright^CD94+NKp46+; stage 5 is CD56^dim^NKp80+Granzyme B+; and stage 6 is CD56^dim^NKp80+Granzyme B+CD57+ (1–7). The developmental trajectory of NK cells is associated with that of helper innate lymphoid cells (ILC) as they both can arise from stage 3 innate lymphoid precursors. In addition, plasticity has been observed in mature ILC and NK cell populations as stage 4 NK cells can also give rise to ILCs (2, 5, 6, 8). The isolation of multipotent NK/ILC progenitors (CD34+integrin β7+) from secondary lymphoid tissues (SLT) that differentiate into mature NK cells in vitro and the spectrum of NK cell developmental stages in tonsil demonstrates that tonsils serve as sites of human innate lymphocyte development (3, 5, 6, 9).

Spatial and functional domains in tonsil include B cell follicles, interfollicular and parafollicular spaces, crypts and subepithelial domains at the exterior of the tissue which lines the oral cavity (10, 11). CD34+CD45RA+ NK cell progenitors and mature NK cells are found in the T cell rich parafollicular domain that surround follicles (9, 12). ILCs and NK cell precursors are located throughout parafollicular and subepithelial domains within SLT, including tonsil (9, 13, 14). Non-immune cells that play roles in SLT organization and function include gp38+ (podoplanin) fibroblastic reticular cells (FRCs), which form a collagen-rich network to support immune cell trafficking through chemokine secretion and physical properties (15). Other non-immune cells include CD31+ endothelial cells and mesenchymal stromal cells (MSCs), which form a heterogeneous stromal cell population, and high endothelial venules (HEVs), which help guide immune cells into tissue via lymphocyte expression of L-selectin and the expression of CCL21, MAdCAM-1, and ICAM-1 on HEV endothelial cells (15–17).

Membrane-bound and secreted factors including IL-3, IL-7, stem cell factor (SCF), FMS-like tyrosine kinase-3 ligand (Flt3L), Delta-like canonical notch ligand-1 (DLL1), and IL-15 are important for inducing NK cell development from CD34+ progenitors and are likely provided by both immune and non-immune cells in sites of development (8, 18, 19). Endothelial cells and MSCs, including FRCs, can also support in vitro NK cell differentiation (20–23). Direct contact between stromal cells and NK progenitors is required to promote NK cell development and proliferation, suggesting that stromal cells are in proximity to NK progenitors in vivo (24–27). Similarly, immune cells in SLT, including T cells and dendritic cells, express ligands and cytokines including IL-2, Flt3L, and IL-15, which can provide maturation signals to developing NK cells (12, 28, 29). The location of developing and mature NK cells is influenced by chemokine receptors and their ligands. CCR7 ligands CCL19 and CCL21 promote homing of mature lymphocytes to SLT and can also recruit common lymphoid progenitors to the thymus to undergo T cell differentiation, suggesting they likely play a role in guiding earliest tissue-seeding precursors from blood to tonsil (30, 31).

T cell differentiation can be mapped by the spatial localization of signals that promote stage-specific localization and differentiation (32). Based on previous studies, our current understanding of NK cell development is that circulating CD34+ progenitors enter SLT through HEVs in the inter-follicular domain and migrate to the parafollicular zone to undergo maturation into functionally mature NK cells (4). However, the spatial localization of NK cell developmental subsets in relation to their trafficking and developmental niche has not been comprehensively defined in human SLT. Here we measure the spatial distribution of mature NK cell subsets, NK/ILC progenitors, and tissue resident NK cells in tonsil using cyclic immunofluorescence (CyCIF) microscopy and imaging mass cytometry (IMC). Quantification of the proximity of NK cell subsets to other immune subsets, stromal cells, and developmentally supportive ligands demonstrates that stages of NK cell development occur at defined sites within tonsil that provide ligands and proliferative signals required to generate mature NK cells. Finally, we show the redistribution of NK cell subsets in the presence of inflammation, thus linking changes in local environment to the changes in localization and proliferation of mature NK cell subsets.

## Results

### Mature NK cell subsets have differential spatial distribution in tonsil

Despite previous descriptions of human tonsil as a site of NK/ILC development, the distribution of mature NK cells and NK cell developmental intermediates relative to other cell subsets within tissue has not been well described. We designed and optimized a panel of 35 markers, plus nuclear staining, to distinguish immune subsets including T cells, B cells, myeloid cells, stromal cells, immature NK populations, and mature NK cell subsets within tonsillar tissue using IMC (33) (Supp. Table 1, Supp. Fig. 1A).

Single cell data was collected and pooled from IMC images following nuclei-based segmentation (Supp. Fig. 1B) of 4-5 domains of interest per donor captured from 5 FFPE pediatric donor sections. Using UMAP based on all markers from all 5 pooled donors, we visualized 102,076 cells and relative expression of our markers of interest (Supp. Fig. 2A-C). Leiden clustering identified 10 cell populations in tonsil (Supp. Fig. 2A). Differences between donors were not major drivers of cluster differences based on UMAP analysis of pooled IMC single cell data (Supp. Fig. 2B). We assigned these 10 clusters as CD8+CD3+ T cells, CD3+ T cells, B cells, NK cells, neutrophils, myeloid cells (macrophage, monocyte, and dendritic cells), stromal cells and unknown cells based on their relative expression of the 35 markers captured plus a nuclear marker (Supp. Fig. 1A, 2C-E). NK cells were identified using CD56, NKp46, NKG2A, and CD94 expression (Supp. Fig. 2D-F).

Leiden clustering based on the 35 markers was performed to further phenotype the NK cell compartment. Using this approach, we identified 9 subsets of mature NK cells in tonsil (Fig. 1A, B; Supp. Fig. 2F). We aligned these to previously described stages of NK cell maturation based on their expression of surface and intracellular markers (1, 2, 4, 6, 9, 34). NK cell clusters 2 (NKp46^high^ stage 5), 4 (GrzmB+ stage 4B), 5 (GrzmA^high^ stage 5), and 8 (GrzmB^high^ stage 5) are characterized by expression of granzyme A or B. Granzyme A or B expression suggests that clusters 2, 4, 5, and 8 are comparable to stage 4B or early stage 5 (CD56^dim^) NK cells in circulation, however low expression of CD16 on these clusters relative to clusters 9 (CD16^high^ stage 5) and 3 (CD57+CD16^high^ stage 6) suggests that clusters 2, 4, 5 and 8 are immature or tissue resident populations (Fig. 1B). Cluster 3 (CD57+CD16^high^ stage 6) appears to be stage 6 NK cells based on their expression of CD57. Due to their high expression of CD16, NK cluster 9 (CD16^high^ stage 5) most resembles circulating stage 5 NK cells. Clusters 1 (stage 4), 6 (CD94^high^ stage 4A-B) and 7 (NKG2A^high^ stage 4B) are reflective of circulating stage 4 NK cells and do not express granzyme B, CD16 or CD57 (Fig. 1B). Cluster 7 represents a stage 4B-like population due to higher expression of NKp80, and cluster 6 represents a seemingly transitional population between stages 4A and 4B based on intermediate NKp80 expression (Fig. 1B). As expected, clusters aligned with stage 4 subsets were the most highly represented when considering number of cells per cluster (Fig. 1B).

**Fig. 1.**
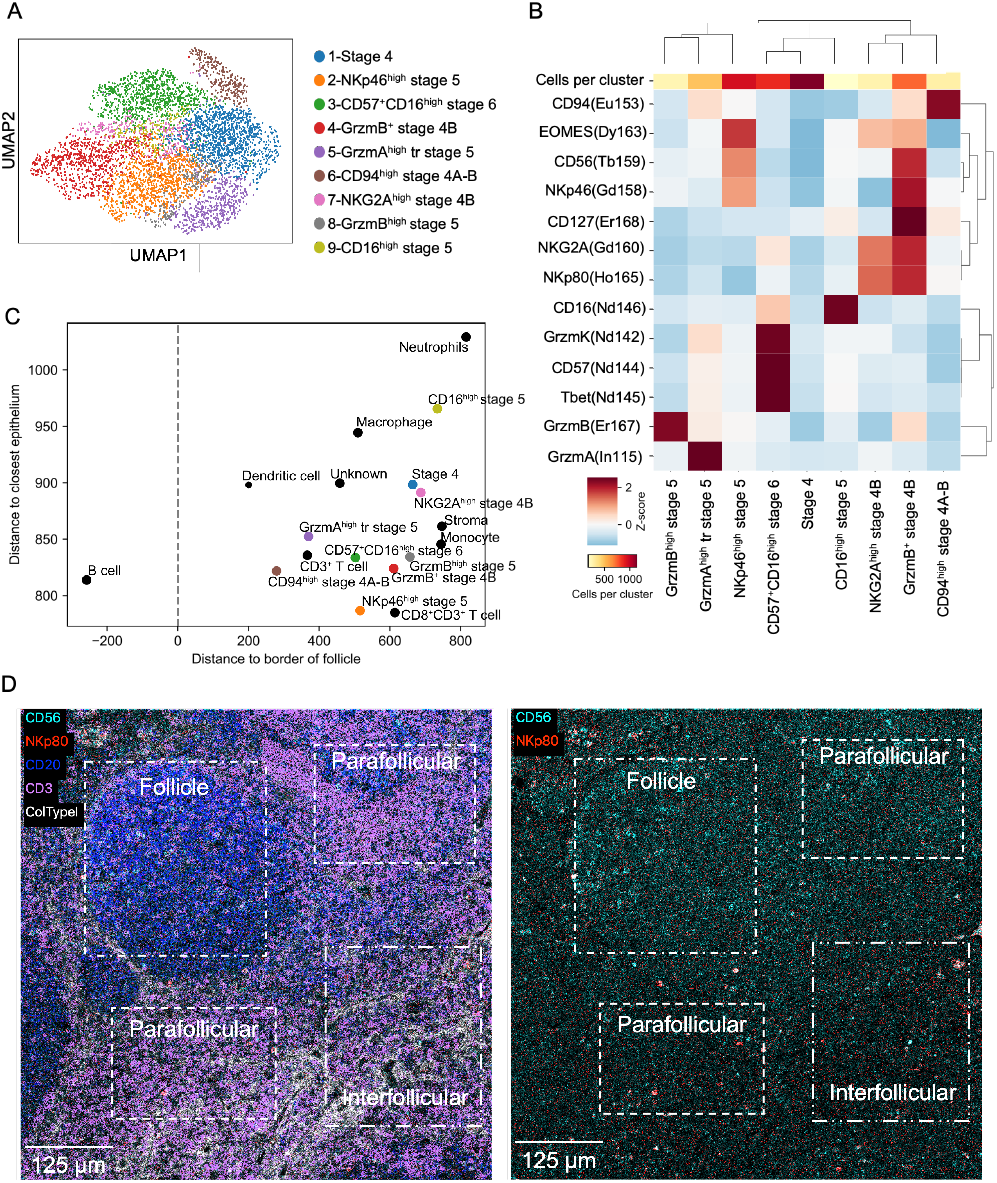
Identification of tonsillar NK cells using imaging mass cytometry (IMC). Pediatric tonsil sections were stained and imaged by IMC. Leiden clustering (0.5 resolution) was performed on NK cells following cell segmentation and primary identification of cell population as described in Methods. A) 9 clusters of NK cells present in tonsil following unsupervised clustering (n=5 donors). B) Relative expression intensity of NK cell markers used to classify NK cell clusters identified by Leiden clustering. C) Average spatial distance plot of NK cell subsets and other tonsil cell populations quantified from the average shortest distance of cells to the nearest epithelium border (y-axis; µm) versus the average shortest distance to the border of the nearest follicle (x-axis; µm). D) Representative image of tonsillar domains [follicles, parafollicular, and interfollicular] taken from an FFPE tonsil section stained with 35 metal-conjugated antibodies plus 191/193-Ir (DNA) and acquired by a Fluidigm Hyperion imaging mass cytometer. Overlaid image (left) shows collagen type-1 (grey), CD56 (cyan), NKp80 (red), CD3 (purple), and CD20 (blue). Overlaid image (right) depicting NKp80 (red) and CD56 (cyan). Scale bar 125 µm. Representative of 31 ROIs from 5 donors.

Having identified NK cell subsets in tissue, we sought to define where within tissue NK cell developmental intermediates were found. Using automated image classifiers detecting CD3, CD20, collagen, vimentin, and pan-keratin, we classified tonsil domains as subepithelial, parafollicular, interfollicular, and follicular and measured the distance of NK cell subsets based on our clustering analysis from these domains (Fig. 1C, D). Measurement of the average shortest distance of individual NK cells and other immune subsets to the nearest epithelium or follicle border was averaged to define the spatial distribution of NK cell subsets in tissue (Fig. 1C). This quantification demonstrated that CD94^high^ stage 4A-B, NKp46^high^ stage 5, GrzmA^high^ tissue resident stage 5, and CD57+CD16^high^ stage 6 NK cells shared similar spatial localization to CD3+ T cells based on their closer proximity (<600 µm) to the nearest follicle border (Fig. 1C). In comparison, stage 4, NKG2A^high^ stage 4B, CD16^high^ stage 5 and GrzmB^high^ stage 5 NK cells were on average >600 µm from the border of the nearest follicle, similar to CD8+CD3+ T cells, monocytes, stroma and neutrophils (Fig. 1C). Stage 4, NKG2A^high^ stage 4B, and CD16^high^ stage 5 were on average >900 µm from the nearest epithelial border, whereas other observed NK cell clusters shared a similar closer distance to follicle borders (Fig. 1C). In agreement with our quantification, visualization of CD56+NKp80+ NK cells showed that most were in the interfollicular and parafollicular domains, with a smaller population present within the follicle and subepithelial domains (Fig. 1D). Together, this shows that most stage 4 and tissue resident-like stage 5 NK cells share a similar spatial localization and are abundant within the parafollicular zone, which is enriched for T cells. In contrast, stage 4, NKG2A^high^ stage 4B, and GZMB+ stage 5 NK cells were found in the interfollicular domain more distal to follicles.

### NK cell progenitors are localized to tonsillar inter- and parafollicular domains

Imaging mass cytometry was not well-suited for the detection of rare cells in complex tissue, as its spatial resolution made it difficult for us to confidently identify single cells in tightly packed domains. With the aim of identifying the earliest NK cell progenitors in tonsil present at low frequencies, we turned to CyCIF microscopy (35, 36) for greater spatial resolution and detection of larger domains of interest. A panel of 68 antibodies (Supp. Table 2) were used in 23 rounds of iterative staining and widefield imaging to detect mature and progenitor NK cell subsets, other immune subsets, and tissue architecture and stromal cells in FFPE sections from 9 pediatric tonsil donors, 3 of which were previously analyzed by IMC. Multiple markers were excluded from subsequent analysis due to lack of specific staining based on their low signal to noise ratio (Supp. Table 2), resulting in a final number of 43 markers. Ki67 was also included in our CyCIF panel to distinguish proliferating from non-proliferating cells. Cell population identities were assigned based on a manual gating strategy which designated a positive/negative threshold for each marker to allow for the detection of rare subsets that would have been overlooked using an unsupervised approach. In addition to other immune subsets, we identified 7 major populations of NK cells (Fig. 2A, Supp. Fig. 3A) that were also detected by flow cytometry (Supp. Fig. 3B) using the gating strategy shown in Supp. Fig. 3A. NK cell subsets included stage 1/2 NK progenitors (‘NK progenitor’, Lin–CD45+CD34+CD122+CD127+CD49d+), stage 4 (Lin–CD45+CD56+CD127+GZMK+/–GZMB–), stage 5 (Lin–CD45+CD56+CD127–GZMB+CD57–) and tissue resident populations (above markers plus CD69+ or CD103+) or proliferating populations (above markers plus Ki67+) (Fig. 2A; Supp. Fig. 3A). The absolute count of each NK cell subset detected in each tonsil domain was enumerated (Fig. 2A; Supp. Fig. 3C) but varied between donors, in part due to differences in ROI size and domains captured in an ROI (Supp. Fig. 3D). Tissue resident and non-tissue resident stage 4 NK cells were the most abundant population of NK cells in all donors, however the frequency of each NK cell subset, including stage 4, varied between donors (Supp. Fig. 3E). We also identified other immune cells including macrophages (CD45+Lin–CD68high), follicular dendritic cells (CD45+Lin–CD11c+CD68–), T cells (CD45+CD3+CD56–), and B cells (CD45+CD20+) (Fig. 2A; Supp. Fig. 3F-H). The inclusion of structural and stromal cell markers allowed for the identification of epithelial cells (CD45– pan-keratin+collagen type 1+) within the subepithelial domain, fibroblastic reticular cells (FRCs) or mesenchymal stromal cells (MSCs; CD45–CD34–CD31–gp38+) which surround and regulate the immune compartments of the tonsil (37, 38), and endothelial cells (CD45–CD34+SMA+CD31+) that make up lymphatic and blood vessels in the interfollicular and parafollicular domains (Fig. 2A, Supp. Figure 3F-H).

**Fig. 2.**
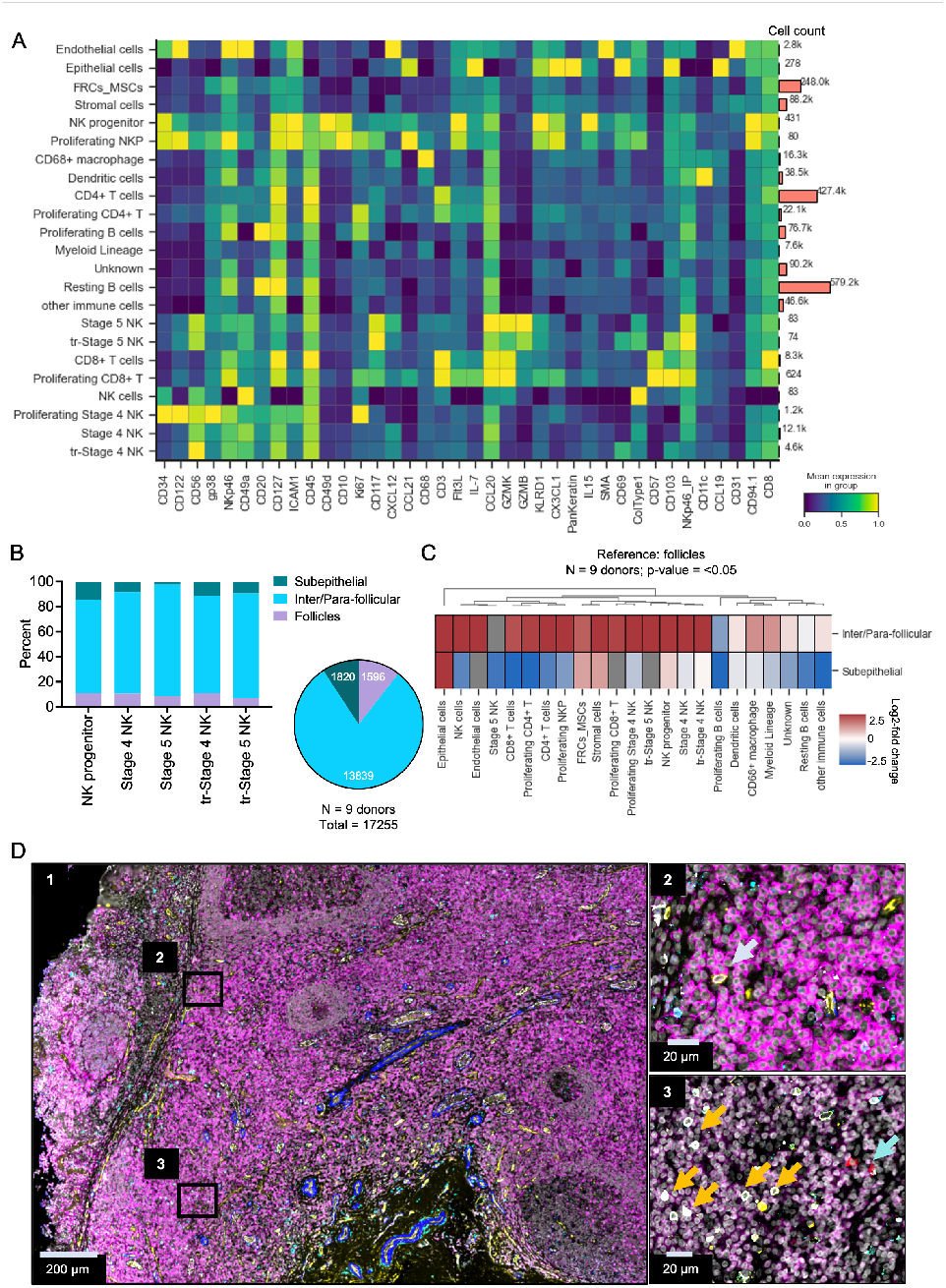
Spatial localization of NK cell subsets in tonsillar tissue revealed by cyclic immunofluorescence (CyCIF). Pediatric tonsil sections were stained and imaged by CyCIF. Immune and non-immune cells, including NK cell subsets, were identified based on their expression of 43 unique markers as described in Methods. A) Heatmap with row clustering and cell count of immune and non-immune cells from pediatric tonsil (n=9 donors; 9 ROIs). B) Frequency of individual (left) and pooled (right) NK cell subsets in subepithelial, follicle, and inter-/para-follicular domains assigned by manual annotation. Frequency was calculated by normalizing the absolute count of each subset in each domain to the total count of cells identified (n=9 donors; 9 ROIs). Domains were manually annotated based on tissue architecture and CD20, CD3, collagen type 1, and pan-keratin staining. C) Log2-fold change in the abundance of NK cell subsets between tonsil domains. Log2-fold change was calculated relative to the frequency of the respective population found within follicle domains; non-significant differences shown in grey (n=9 donors; 9 ROIs). D) Visualization of NK cell subset spatial distribution in tonsil. Representative CyCIF image of FFPE tonsil tissue stained and imaged with 43 markers over 23 cycles. Image 1 is a representative ROI captured by CyCIF and shows an overlay of smooth muscle actin (blue), CD56 (cyan), CD45 (magenta), CD34 (yellow), and CD31 (white). Image 2 is an enlarged image of a CD34+ NK cell progenitor (grey arrow) in the interfollicular domain and depicts the same markers as image 1. Image 3 is an enlarged image of mature stage 4 (orange arrows) and stage 5 (blue arrows) NK cell subsets in the parafollicular domain and depicts GrzmK (yellow), CD56 (cyan), CD3 (purple), and GrzmB (red). Representative of 9 ROIs from 9 donors. Scale bar 200 µm and 20 µm.

To determine the localization of NK cell subsets, we manually annotated tonsil domains, including subepithelial, inter-/para-follicular, and follicular domains, using 9 ROIs from 9 donors. Tonsil domains were identified and annotated based on distribution of type 1 collagen, pan-keratin, CD20, and CD3 (Supp. Fig. 3I). The presence of each NK cell population within all annotated domains was then quantified (Fig. 2B; Supp. Fig. 3C). Our previous findings from IMC analysis, namely that mature NK cells were predominantly in the inter- and parafollicular domains relative to follicle or subepithelial domains, were confirmed by CyCIF (Fig. 2B). We additionally found that >70% of early NK cell progenitors (Lin–CD45+CD34+CD122+CD127+CD49d+) were found within the inter- and parafollicular domains of the tonsil (Fig. 2B). The greater representation of NK cell progenitors in interfollicular and parafollicular space was confirmed by quantifying the log2-fold difference of NK cells in each domain relative to the percent found in follicles and normalizing for the total cells detected to account for difference in domain area (Fig. 2C). This analysis showed that most NK cell subsets, including NK cell progenitors, were more highly abundant in the inter/para-follicular domain relative to the follicle (Fig. 2C). NK cell progenitors could be visualized deep within the interfollicular and parafollicular domains near blood vessels (Fig. 2D, image 2). Tissue resident and non-tissue resident stage 4 NK cell populations were the most abundant NK cell populations in each domain (Fig. 2B, C), and preferentially localized to the inter/para-follicular domain relative to follicular or subepithelial domains (Fig. 2B, C and Supp. Fig. 3C). Stage 5 NK cells were not highly detected in subepithelial domains (<2%) (Fig. 2B) but were enriched in the inter/para-follicular domains (Fig. 2C). As previously described, we observed stage 4 (orange arrow) and stage 5 (blue arrow) NK cells within the inter- and parafollicular domains (12) (Fig. 2D). Therefore, with higher resolution imaging we were able to detect the spatial localization of rare NK cell progenitors (grey arrow; Fig. 2D) and further validate the distribution of mature NK cell subsets. NK cell progenitors, comprising <10% of the total NK cell population (Supp. Fig. 3E), were enriched within the inter/para-follicular zone but were also found in the follicle and subepithelial domains. Non-tissue resident and tissue resident stage 4 and stage 5 NK cells were also primarily enriched to inter/para-follicular or follicle domains.

### Stage-specific changes in spatial distribution of NK cell developmental subsets relative to other tonsil cell populations

Having quantified the spatial localization of NK cell subsets, we sought to better understand their interactions with other immune subsets. Using cell masks colored by assigned cell identities from a representative CyCIF image, we observed that NK cells subsets engaged in unique interactions dependent on their developmental stage and tissue location (Fig. 3A). Specifically, stage 4 NK cells engaged with many cell types including T cells, stromal cells, and B cells, while stage 5 NK cells appeared to interact primarily with T cell populations in interfollicular and parafollicular domains (Fig. 3A). Tissue resident stage 4 NK cells (white arrows) were adjacent to stromal cells in the interfollicular, parafollicular, and subepithelial space (Fig. 3A).

**Fig. 3.**
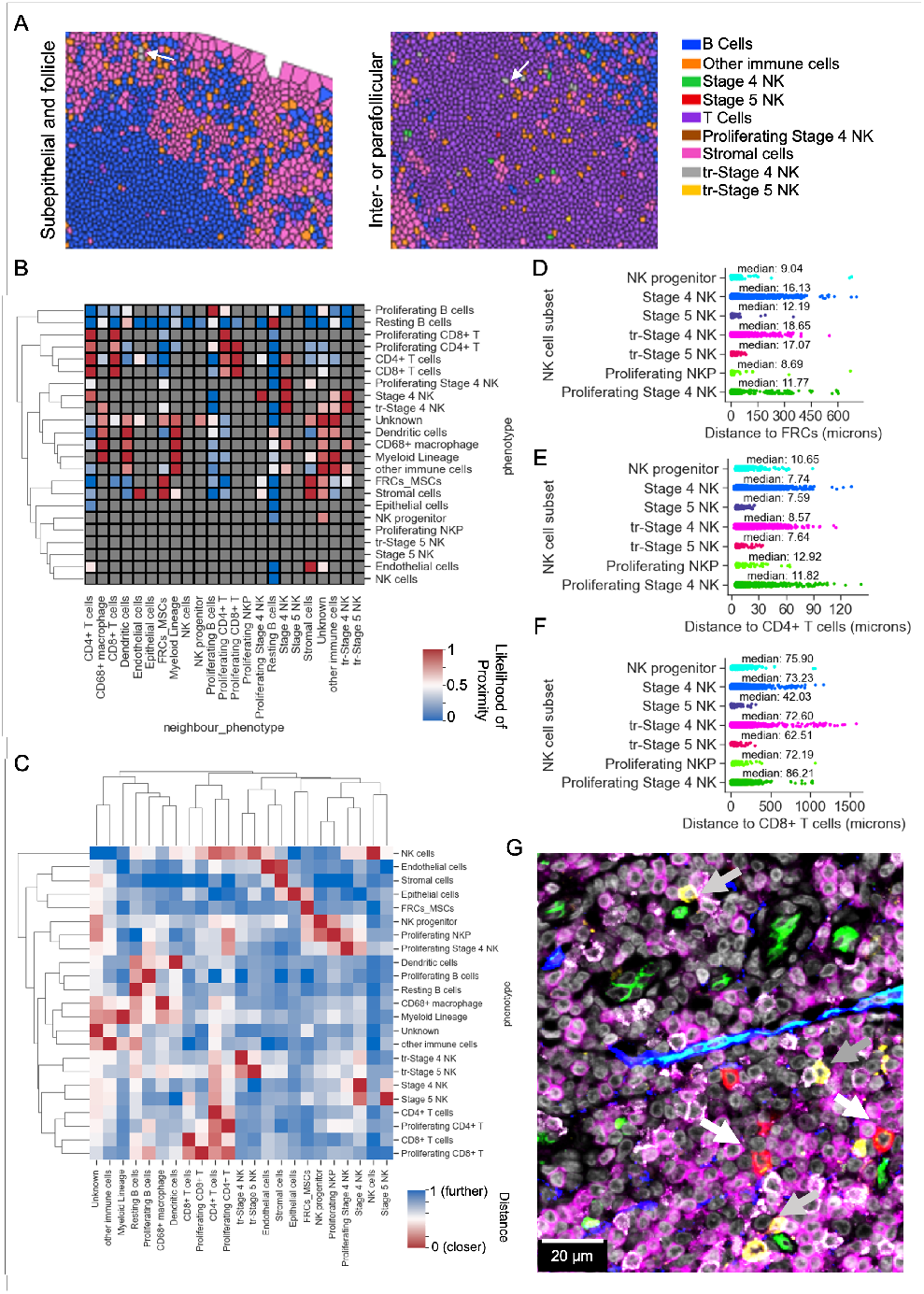
NK cell interactions in tonsillar tissue change through development. CyCIF images were segmented and immune and non-immune cells, including NK cell subsets, were identified based on their expression of 43 unique markers. A) Representative cell mask showing distribution of tonsil cell populations identified by CyCIF. Cell mask was derived from Mesmer nuclear segmentation mask of CyCIF ROIs and colored by assigned cell identities using SCIMAP Voronoi plot function as described in Methods. B) Likelihood of cell-cell interactions between tonsil cell populations identified by SCIMAP cell co-occurrence analysis function with a radius of 22.7 µm (50 pixels). Heatmap shows significant increased likelihood as red, decreased likelihood as blue, and non-significant neighbors represented by grey (mean of n = 9 donors; 9 ROIs). C) Heatmap, with row and column clustering, of average shortest distance between tonsil cell populations in microns. Average shortest distance was calculated for all phenotypes by taking the mean of the shortest distance between a cell of interest and the nearest cell of a given population of interest (n = 9 donors; 9 ROIs). A smaller average shortest distance is represented by red, and a larger average shortest distance is represented by blue. D) Distribution of the median shortest distance between NK cell subsets and gp38+ FRCs (n=9 donors; 9 ROIs). E) Median shortest distance between NK cell subsets and CD4+ T cells (n=9 donors; 9 ROIs). F) Median shortest distance between NK cell subsets and CD8+ T cells (n=9 donors; 9 ROIs). G) Representative CyCIF image of FFPE tonsil section zoomed in on non-tissue resident stage 4 (grey arrows) and stage 5 (white arrows) NK cells interacting with CD4+ and CD8+ T cells based on CD3 (magenta), CD56 (yellow), GrzmB (red), CD34 (green), and CD8 (white). Representative of 9 ROIs from 9 donors. Scale bar 20 um.

To quantify these observations, we performed radius spatial co-occurrence analysis using SCIMAP (39) to measure the likelihood of proximity between two cell populations. We first identified the neighbors to every cell within a 50-pixel (22.7 µm) radius from the center of each cell and computed significant recurring interactions to define subsets of cells found in the vicinity of NK cell subsets (Fig. 3B). We found that stage 4 NK cells were in areas enriched for CD4+ T cells, and not found in domains significantly enriched for proliferating (Ki67+) B cells (Fig. 3B). This observation is reflective of the spatial localization of stage 4 NK cells (Fig. 2) and suggests increased propensity for more mature NK cells to be in the T cell rich zone of the parafollicular domain (Fig. 3B). When considering tissue residency, we found that tissue resident stage 4 NK cell populations had a unique niche and were found primarily near CD68+ macrophages and other NK cell subsets, but not B cells (Fig. 3B). Notably, non-tissue resident NK cells and tissue resident stage 5 NK cells were not found in significant proximity to any other cell population identified by CyCIF (Fig. 3B). Due to the rarity of proliferating NK progenitors, we did not detect significant enrichment of other identifiable cell populations in their proximity using this population-based approach, however NK progenitors were unlikely to be found in the same domains as resting (Ki67–) B cells (Fig. 3B).

Next, we quantified the likelihood of cell-cell interactions of NK cell subsets by measuring individual cell coordinates and calculating the average shortest distance (µm) between two cell populations of interest to generate a relative distance. This approach complements neighborhood analysis by measuring the shortest average distance between cell populations to define their relative proximity. We observed that more mature NK cell populations (stage 4 and 5) exhibited a higher tendency to be in proximity to other NK cells (Fig. 3C). We also confirmed that tissue resident stage 4 and 5 both were in closer proximity to myeloid cells including CD68+ macrophages than non-tissue resident NK cells (Fig. 3C). When comparing the distance between cell populations we found that non-proliferating (Ki67–) and proliferating (Ki67+) NK cell progenitors were uniquely situated closer to FRCs relative to mature non-tissue resident NK cell populations (Fig. 3C). Measuring the distribution of the distance of NK progenitors to FRCs confirmed this finding and demonstrated that NK progenitors were uniformly closer in distance (mean = 18.72 µm, median = 9.04 µm) to FRCs than mature (stages 4) NK cells (Fig. 3D). We also observed a higher frequency of stage 4-5 NK cells more closely in contact with CD4+ T cells in comparison to NK cell progenitors (Fig. 3C, E). Stage 5 NK cells were in closer proximity (median = 42.03 µm) to CD8+ T cells than stage 4 (median = 73.23 µm) and progenitor cells (median = 75.90 µm; Fig. 3C, F). Similar observations were made for tissue resident populations proximity to CD4+ and CD8+ T cells (Fig. 3C, E, F). Consistent with these measurements, we observed stage 4 and stage 5 in proximity to CD4+ and CD8+ T cells, respectively, within the parafollicular region (Fig. 3G). Overall, NK cell subset interactions not only change with maturation and tissue residency but are also reflective of their spatial localization. Based on neighborhood analysis, NK progenitors occupy different domains primarily adjacent to stromal cells within the interfollicular and parafollicular domains. In contrast, stage 4 and 5 NK cells are more localized to T cell rich zones.

### NK progenitors proliferate adjacent to stromal cells in tonsil

As tissue seeding precursors are rare in tonsil but can generate robust NK cell developmental intermediate and mature NK cell populations, we sought to further define the relationship between NK progenitors and cell proliferation. NK progenitors were more proliferative than stage 4 NK cells as seen by CyCIF (Fig. 4A). We confirmed these findings by flow cytometry of tonsil NK cells from 7 pediatric donors, which identified stages 1 and 2 NK cells with significantly higher expression of Ki67 than stages 3-5 (Fig. 4B). Quantification of the localization of proliferating cells demonstrated that proliferating NK progenitors and stage 4 NK cells were primarily found in inter- or parafollicular domains, with some proliferation in follicles and subepithelial spaces (Fig. 3C, Fig. 4C). Since most NK cells were proliferating within the inter- and parafollicular domains (Fig. 4C) we sought to define the interactions that may influence NK cell proliferation. We measured the distance of proliferating subsets to FRCs, as stromal cells can present cytokines to lymphocytes in primary and secondary lymphoid tissues (26, 40–42). Radius spatial co-occurrence analysis first revealed that proliferating (Ki67+) stage 4 and 5 NK cells were frequently found near gp38+ FRCs (Fig. 3B). Quantification of the shortest average distance of proliferating subsets to each cell type confirmed differences in interactions with FRCs (Fig. 3C, 4D). Proliferating (Ki67+) and non-proliferating NK progenitors appeared to consistently be near FRCs within tonsil (Fig. 3C-D). Stage 4 NK cells were more likely to be in contact with gp38+ FRCs and MSCs than non-proliferating or tissue resident stage 4-5 NK cell populations (Fig. 3B-D, 4D-E). We also saw an increase in the average shortest distance between CD4+ T cells and subsets of proliferating stage 4 NK cells (Fig. 3B, C, E). We also observed close interactions between gp38+ FRCs and proliferating stage 4 NK cells based on gp38, Ki67, CD56, and granzyme B staining (Fig. 4D).

**Fig. 4.**
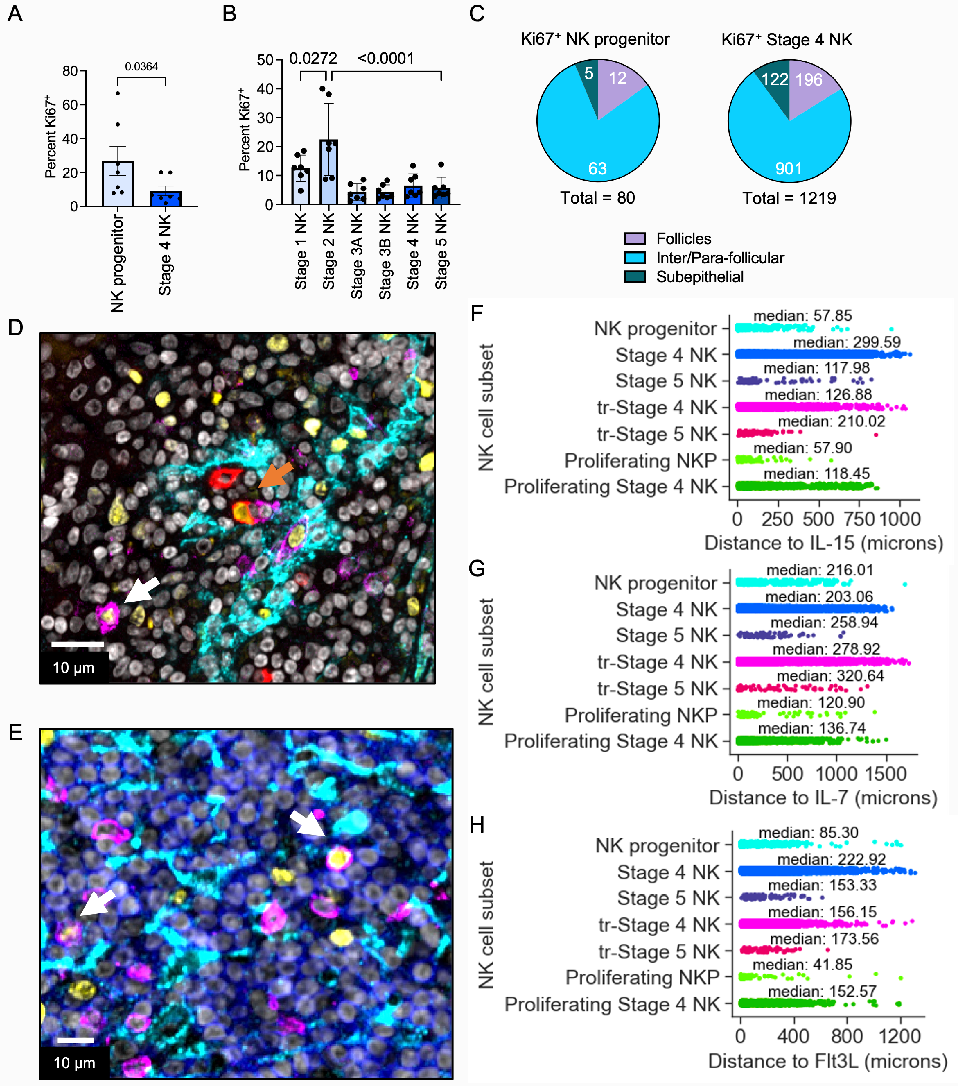
Human NK cell progenitors proliferate and localize to cytokine expressing cells and gp38+ fibroblastic reticular cells. A) Percent of progenitor, stage 4, and stage 5 NK cells expressing Ki67 quantified from CyCIF single cell data (n=9 donors; 9 ROIs). Ki67 positive NK cells were identified by manual gating based on Ki67 expression intensity. Mean and standard error were plotted; p-value was determined by multiple unpaired T-test. B) Percent of Ki67+ stage 1-6 NK cell subsets quantified by flow cytometry. Tonsil mononuclear cells were isolated from 7 donors and stained with the antibody panel described in Supp. Table 3 and subsets were identified using the gating strategy in Supp. Fig. 2A and B. Mean and standard error were plotted; p-values were determined by one-way ANOVA with multiple comparisons. C) Spatial distribution of proliferating (Ki67+) NK cell subsets. Percentage of proliferating NK cells in each tonsil domain was quantified by dividing the number of Ki67+ NK cells in each domain by the absolute count of the respective population (n = 9 donors; 9 ROIs). D) Representative CyCIF image of proliferating stage 4 (white arrow) and stage 5 (orange arrow) NK cells in proximity to gp38+ FRCs within the interfollicular domain. Image depicts a cropped ROI from a larger field of view of a FFPE tonsil section. CD56 (magenta), Ki67 (yellow), GrzmB (red), gp38 (cyan), and DAPI (grey). Scale bar 10 um. E) Representative image of proliferating NK cells adjacent to T cells and gp38+ FRCs within the parafollicular domain. Image is a cropped ROI from a larger field of view of a FFPE tonsil section. CD56 (magenta), Ki67 (yellow), CD3 (blue), gp38 (cyan), and DAPI (grey). Scale bar 10 um. F-H) Mean shortest distance between proliferating NK cell subsets and cytokine expressing cells including (F) IL-15+, (G) IL-7+, and (H) Flt3L+ cells. Cells positive for IL-15, IL-7, and Flt3L were identified by manual gating of cell populations identified by CyCIF and distanced was measured as described in Methods (n=9 donors; 9 ROIs).

NK cell proliferation and maturation are dependent on cytokine presentation by developmentally supportive cells and other immune cells. Quantification of the median shortest distance between NK cell subsets was used to characterize interactions between non-proliferating and proliferating NK cell subsets and cytokine producing cells including IL-15+, IL-7+, or Flt3L+ cells (Fig. 4F-H). Considering the median shortest distance of proliferating NK cell populations to IL-15 expressing cells, subsets of Ki67+ stage 4 NK cells were closer than non-proliferating stage 4 NK cells to IL-15+ cells (Fig. 4F). Interestingly, regardless of Ki67 expression, NK cell progenitors appeared to be in closer proximity to IL-15+ cells than proliferating and non-proliferating stage 4 and 5 NK cells (Fig. 4F). Tissue resident NK cells were closer to IL-15+ cells (Fig. 4F) than non-tissue resident, and proliferating, stage 4 and 5 NK cells.

Relative to IL-15, proliferating (Ki67+) NK progenitors were farther from IL-7 expressing cells but were the closest subset to them, and stage 4 NK cells were less likely to be in proximity to IL-7+ cells compared to IL-15+ cells (Fig. 4G). Although proliferating progenitor and stage 4 NK cell subsets did not appear to be as close to IL-7 expressing cells than they are to IL-15, proliferating subsets were in closer proximity to IL-7+ cells than non-proliferating subsets (Fig. 4G). Non-proliferating (Ki67–) NK cell subsets all had a median distance of >200 µm to the nearest IL-7+ cell (Fig. 4G).

Next, we measured the shortest average distance of NK cell subsets to Flt3L+ cells. Proliferating (Ki67+) NK cell progenitors were the closest in proximity to Flt3L+ cells in comparison to more mature stage 4 proliferating subset (Fig. 4H). Proliferating NK cell progenitors had a median distance of approximately half (median = 41.85 µm) that of non-proliferating NK cell progenitors (median = 85.3 µm; Fig. 4H). NK cell subsets were in closer proximity to Flt3L+ cells when proliferating regardless of developmental stage (Fig. 4H). Tissue resident stage 4 and 5 NK cells were also seen to be closer to Flt3L+ cells than non-tissue resident populations (Fig. 4H). Furthermore, our data shows that there is a spatial relationship between FRCs and proliferating NK cell subsets and cytokines, which suggests that FRCs can play a role in establishing a developmental or proliferative niche for NK cells in tonsil. As predicted by their well-described roles in NK cell development, presentation of IL-15 is associated with NK cell maturation and proliferation in tissue, and IL-7 and Flt3L expression are associated with NK cell progenitor proliferation.

### Chemokine receptor expression and chemokine lig- and localization in tonsil

Based on the distribution of NK cell subsets in tonsil, we predicted that their localization could be dependent on differential chemokine receptor expression on NK cell subsets and chemokine ligand expression in tonsil domains. We used previously published RNA-seq data from sorted NK cell developmental subsets to define the relative gene expression of selected chemokine receptors on NK cell subsets (stages 3-5) from pediatric tonsil (43) (Fig. 5A, Supp. Fig. 4A). We found that tonsillar stage 3 and 4A NK cells have higher expression of CXCR6, CCR6, CCR4, and CXCR5 relative to later stages of NK cell development, whereas tonsillar stage 4B and 5 NK cells have increased CCR1, CCR5 and CXCR3 gene expression (Fig. 5A, Supp. Fig. 4A). CX3CR1 is upregulated in tonsillar stage 5 NK cells and CXCR4 was highly detected at the transcript level in stages 3-5 tonsillar NK cells relative to other chemokine receptors (Supp. Fig. 4A), with higher expression in stages 3 and 5 than stage 4 (Fig. 5A). Together, these data identified patterns of chemokine receptor expression that generally differed between intermediate NK cell progenitors (stages 3 and 4A) and more mature NK cells (stages 4B and 5).

**Fig. 5.**
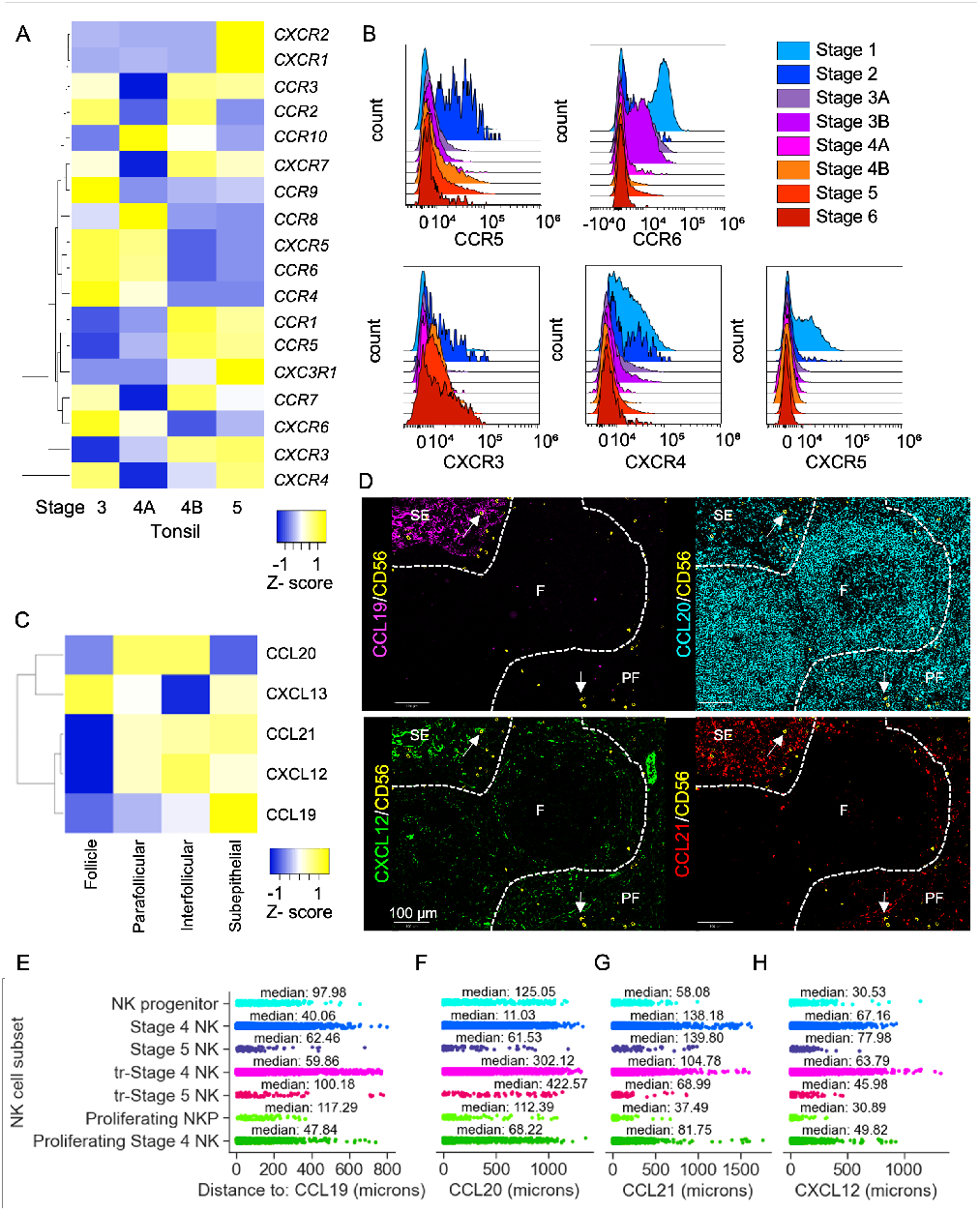
Chemokine receptor expression through NK cell development changes with maturation and influences localization of NK cells in tonsil tissue. A) Normalized gene expression (row scaled FPKM) from bulk RNA sequencing of sorted NK cell developmental subsets from 12 tonsil donors pooled into 3 replicates (Hegewisch-Solloa et al., 2021). B) Flow cytometric analysis of NK cell chemokine receptor protein expression. Mononuclear cells were isolated from disassociated tonsil tissue and stained with 1 of 4 unique panels outlined in Supp. Table 4 (6-7 donors per panel). C) Average fluorescence intensity of CCL19, CCL20, CCL21, CXCL12, and CXCL13 in tonsil microdomains. Microdomains were annotated using HALO AI’s trained network. Calculated from 3 ROIs from 3 donors. D) Representative CyCIF image of FFPE tonsil section showing distribution of CD56+ cells (yellow; white arrows) and CXCL12 (green) CCL19 (magenta), CCL20 (cyan), or CCL21 (red). Image depicts three microdomains including subepithelial (SE), follicle (F), and parafollicular (PF) domains; interfollicular domain was not included due to the similar chemokine ligand expression as parafollicular domain. E-G) Median shortest distance (microns) NK cell subsets to (E) CCL19, (F) CCL20, (G) CCL21, or (H) CXCL12 expressing cells in tonsil (n=9 donors; 9 ROIs). Distance (x-axis) was calculated for each cell of a given population of interest to the nearest CCL19+, CCL20+, CCL21+, or CXCL12+ cell.

To further define chemokine receptor protein expression on NK cells and progenitors, paired peripheral blood and tonsil samples from 6-7 pediatric donors were stained with 4 19-marker flow cytometry panels to identify NK cell developmental subsets and measure their expression of 12 chemokine receptors (Supp. Table 4). Stage 1 NK cells (Lin–CD45+CD34+CD117–; lineage markers = CD3, CD19, and CD14) from tonsil expressed higher levels of CXCR4 and CXCR5 relative to more mature stages of tonsil NK cells (Fig. 5B). CCR6 is expressed by tonsil stage 1 NK cells and is detected up to stage 3B, and the transition from stage 3B to stage 4A is marked by downregulation of CCR6. Stage 2 NK cells in tonsil, but not peripheral blood, uniquely express CCR5 (Fig. 5B; Supp. Fig. 5B). Stages 4B-6 have increased expression of CXCR3 relative to stages 1-4A (Fig. 5B). A subset of tonsil stage 6 NK cells expresses CX3CR1, but it is not detected on early stages except on a subset of stage 1 and 2 (Supp. Fig. 5A). While in peripheral blood NK cell subsets, CX3CR1 and CXCR3 is more highly detected on stage 4-6 (Supp. Fig. 5B). When comparing expression of chemokine receptors between tonsil and peripheral blood NK cell subsets by flow cytometry, we also noted differences in CCR4, CCR5, CCR6, CXCR4, CXCR5, and CXCR6 (Supp. Fig. 5A, B). CCR4 is expressed by a subset of peripheral blood stage 1 NK cells but not on tonsil stage 1 cells (Supp. Fig. 5A, B). In contrast, CCR6 is expressed by a population of tonsil stage 1 NK cells, while absent on peripheral blood stage 1 NK cells. In addition, a subset of tonsil stage 1 NK cells expresses CXCR5, which is not reflected in peripheral blood. We did not detect robust levels of CCR1, CCR4, CCR7, CCR8, and CCR9 by flow cytometry on tonsil NK cell subsets, although they are detected by RNA-seq (Supp. Fig. 4A, 5A). CXCR1, CXCR2, CXCR7, CCR3, and CCR10 were also not detected at high levels [normalized FPKM <1.5] in tonsil NK cell subsets by bulk RNA-seq (Supp. Fig. 4A).

To define the expression of chemokine ligands in tonsil that can interact with NK cell developmental subsets, we performed CyCIF microscopy and quantified the relative expression of chemokine ligands in subepithelial, interfollicular, parafollicular, and follicular domains (Fig.5C, D). Follicular domains were marked by high expression of CXCL13 (CXCR3 and CXCR5 ligand), while CCL19 (CCR7 and CCR11 ligand) was highly expressed in subepithelial domains and CCL21 (CCR7 and CCR11 ligand) was more widely expressed in inter- and parafollicular domains (Fig. 5C). Cells expressing CXCL12 (CXCR4 ligand) were in the parafollicular and interfollicular domains, and to a lesser extent in subepithelial domains (Fig. 5C). Imaging of CXCL12 defined its expression specifically on HEVs and stroma, with mature CD56+ NK cells in proximity to CXCL12+ cells (Fig. 5D). Imaging of CCL19, CCL20, and CCL21 confirmed the less restricted expression of CCL21 than CCL19 and expression of CCL20 in follicles (Fig. 5D).

We performed proximity analysis to measure the shortest average distance from NK cells to chemokine ligand expressing cells. On average, non-tissue resident stage 4 and 5 NK cells, regardless of Ki67 expression, were in closer proximity to CCL19 expressing cells than NK progenitors and tissue resident subsets (Fig. 5E). The distance of stage 4 NK cells to CCL19+ cells was noted to be more heterogenous than progenitor and stage 5 NK cell subsets, for which the majority were <400 µm from CCL19+ cells yet stage 4 NK cells had a lower median distance to CCL19+ cells (Fig. 5E). When measuring the proximity of NK cell subsets to CCL20+ cells we found that tissue resident stage 4 and 5 NK cells were the least likely to be near CCL20+ cells as they had a median distance of 302.12 µm and 422.57 µm respectively (Fig. 5F). Compared to tissue resident and progenitor populations, more mature non tissue resident stage 4 and 5 populations were more likely to be near CCL20+ cells with a median distance of 11.03 µm and 61.53 µm respectively (Fig. 5F). Tissue resident stage 4 and 5 NK cells both shared a median distance of >300 µm to CCL20+ cells suggesting their localization is not related to CCL20 (Fig 5F). Proliferating (Ki67+) stage 4 NK cells were not as likely to be near CCL20+ cells with a median distance of 68.22 µm than non-proliferating stage 4 NK cells yet were closer than proliferating (Ki67+) NK progenitors which had a median distance of 112.39 µm to CCL20+ cells (Fig. 5F). Together this suggests that as NK cell mature from CD34+ multipotent progenitors to stage 5 NK cells, they move closer to CCL19+ and CCL20+ cells in tonsil.

As CCL21 marked the parafollicular, interfollicular, and subepithelial domains of the tonsil (Fig. 5C), we hypothesized that NK cell subsets would be near CCL21+ cells in tonsil. NK progenitor cells were observed to have the shortest median distance (58.08 µm) to CCL21+ cells (Fig. 5G). Proliferating (Ki67+) NK progenitors appeared to be even closer than non-proliferating NK progenitors to CCL21+ cells with a shortest median distance of 37.49 µm (Fig. 5G). Meanwhile, more mature non-tissue resident populations did not appear to be proximal to CCL21+ cells, as both stage 4 and 5 had a median distance of approximately 140 µm to CCL21+ cells (Fig. 5G). Compared to non-tissue resident, tissue resident NK cell populations were closer to CCL21+ cells, yet tissue resident stage 5 NK cells (median = 68.99 µm) were closer to CCL21+ cells than stage 4 (Fig. 5G). Proliferating (Ki67+) NK cell subsets, including progenitor and stage 4, were closer to CCL21+ cells than non-proliferating (Ki67–) subsets (Fig. 5G). Unlike the observations of median distance to CCL19 and CCL20, NK cell subsets appeared to be increasingly further away from CCL21+ cells as they matured. NK cell progenitors, which express higher CXCR4 than more mature NK cells (Fig. 5B), had the closest proximity (median = 30.53 µm) to CXCL12+ cells than more mature NK cell subsets (Fig. 5H). Non-tissue resident and tissue resident stage 4 NK cells shared a similar proximity to CXCL12+ but was not the case for stage 5 subsets (Fig. 5H). Tissue resident stage 5 NK cells were found to have a 30 µm difference in median distance to CXCL12 expressing cells when compared to non-tissue resident stage 5 NK cells (Fig. 5H). Ki67 expression did not influence NK progenitor proximity to CXCL12+ cells, but more mature Ki67+ stage 4 NK cells were uniformly closer to CXCL12+ cells than non-proliferating (Ki67–), non-tissue resident populations by about 15 µm (Fig. 5H). Based on these observations, NK cell progenitors and proliferating subsets localize to CXCL12 in tonsil and as NK cells mature, they move away from CXCL12+ (Fig. 5H).

Overall, our data reveals that progenitor NK cell localization is related to CCL21 and CXCL12 spatial distribution in tonsil. Similarly, we found that proliferating and tissue resident NK cell subsets were more closely localized to CCL21 and CXCL12 expressing cells than non-tissue resident NK cell subsets. Spatial analysis showed that non-tissue resident NK cells are heterogeneously distributed with respect to CCL19, CCL20, CCL21, and CXCL12.

### Identification of NK cell developmental niches in tonsil

In addition to chemokines, we sought to define niches that support NK cell development within human tonsil using CyCIF, Nanostring spatial transcriptomics, and flow cytometry. Nanostring GeoMx transcriptomics analysis of 32 manually defined ROIs from pediatric tonsil sections from 6 donors revealed that parafollicular and interfollicular domains had generally higher expression of developmental cytokines and ligands than follicles (Fig. 6A). Specifically, we found that *DLL4, FLT3L, SCF, IL7, JAG1* and *JAG2* transcripts were more highly expressed by cells within interfollicular and/or parafollicular domains (Fig. 6A). IL-15 did not appear to be highly expressed in tonsil although we detected higher level of transcripts in the interfollicular and parafollicular domains (Fig. 6A). We visually confirmed the differential expression of these ligands by immunofluorescence, which demonstrated the expression of IL-7 in the parafollicular and subepithelial domains and IL-15 and Flt3L in inter-follicular domains (Fig. 6B). As expected, the expression of these ligands was largely on stromal and endothelial cells, and we found that gp38+ FRCs expressed IL-7 and IL-15 in the interfollicular, parafollicular, and subepithelial domains (Fig. 6B). Expression of IL-15 was also detected in the follicle but was not colocalized with gp38 staining, suggesting another source of IL-15 exists in the follicle other than FRCs (Fig. 6B).

**Fig. 6.**
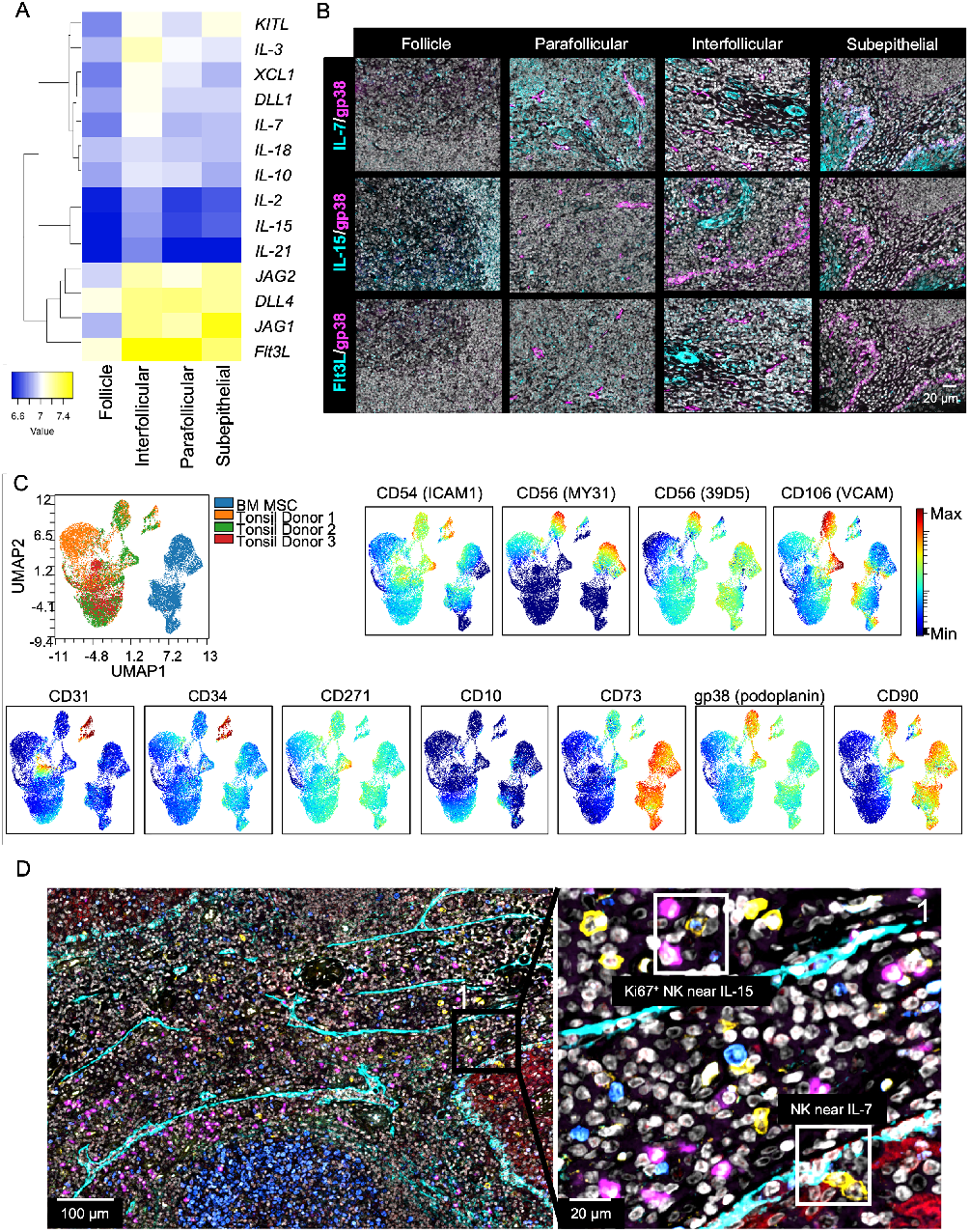
Tonsils harbor cytokine producing stromal cells that provide a developmental niche which supports NK cell maturation and proliferation. A) Nanostring GeoMx transcriptomic analysis of Notch ligands, cytokines, and chemokines known to have a role in NK cell development. Nanostring GeoMx was performed on 32 ROIs from 6 tonsil donors and gene expression data was normalized and log scaled. B) Representative CyCIF image of IL-7 (top panel; cyan), IL-15 (middle panel; cyan), and Fl3tL (bottom panel; cyan) shown in cyan and overlaid with gp38 as a marker for FRCs and MSCs. Scale bar 20 um. C) UMAP projection of tonsil (n=3 donors) and bone marrow stromal cells (n=1 donor) showing expression intensity of endothelial and mesenchymal stromal cell markers. Tonsil and bone marrow stromal cells were isolated and stained with antibody panel in Supp. Table 5 for flow cytometric analysis. UMAP projection was calculated using expression data of all markers except for lineage markers (CD45, CD3, CD14, CD19) and live cell marker. D) Proliferating NK cells in proximity to IL-15 and IL-7 expressing cells including gp38+ FRCs in the interfollicular domain. Representative CyCIF image of IL-15 (magenta), IL-7 (red), gp38 (cyan), CD56 (yellow), Ki67 (blue), and DAPI (grey) staining from FFPE tonsil section. Right image is an enlarged region of interest from the image on the left. Representative of 9 ROIs from 9 donors. Scale bar 100 µm (left) and 20 µm (right).

To better characterize stromal cell subsets in human tonsil and compare them to bone marrow stromal cells that can promote NK cell development, we used a flow cytometry panel designed to measure expression of adhesion ligands (ICAM-1, VCAM-1) and markers of mesenchymal (CD90, CD73) or endothelial state (CD31, CD34) (Supp. Table 5) (15, 25, 26, 37, 38, 44). We additionally included NCAM (CD56), given its previous implication in facilitating human NK cell differentiation (45). Single-cell analysis revealed that bone marrow and tonsil stromal cells clustered distinctly when visualized by UMAP and clearly identified a population of tonsil FRCs highly expressing gp38 and CD73 (Fig. 6C). Markers including CD73 and CD90 were expressed on bone marrow stroma and on a subset of tonsil stroma, whereas ICAM-1 (CD54) and VCAM-1 (CD106) were more highly expressed on a subset of tonsil stroma than bone marrow stroma. As previously described, a clone of CD56/NCAM (39D5) that specifically recognizes bone marrow stromal cells was detected on a subset of both bone marrow and tonsil stroma, as was conventionally detected CD56/NCAM (MY31) (Fig. 6C). The proximity of NK cells to cytokines and ligands on FRCs (Fig. 5) suggests that FRCs are a primary source of proliferative signals for NK cell subsets in tonsil, and visualization of proliferating NK cells adjacent to these stromal cell populations further confirms this observation (Fig. 6D). Further, similarities between subsets of tonsil and bone marrow stroma suggest that analogous populations may act at each site to help promote NK progenitor survival and expansion (46).

### Inflammation changes NK cell subset frequency, cell interactions and localization

Lastly, we aimed to define the effects of inflammation on NK cell developmental subset frequency, cell-cell interactions, and localization. Inflammation in tonsils used for our study was determined by post-operative pathology reports, which determined the presence of chronic or acute local inflammation at the time of resection (Supp. Table 6). We additionally validated donor pathology reports using spatial transcriptomics to evaluate trends in *IFNG, TNF*, and *NFKB1* gene expression. Although *IFNG* expression was not different between noninflamed and inflamed tonsils, *TNF* and *NFKB1* expression was elevated in samples from donors with reports of chronic or acute local inflammation (Supp. Fig. 6A).

Using CyCIF to assess changes in NK cell subset frequency, we found that non-tissue resident stage 4 NK cells make up a higher percentage of the total NK cell population in tonsil in the presence of local inflammation (Fig. 7A). This observation was also reflected in the absolute number of NK cells detected in each tonsil domain from noninflamed and inflamed donors (Supp. Fig. 6C). We calculated the log2-fold difference of NK cell subset abundance using non-inflamed donors as the reference and normalizing cell abundance to the total number of cells detected within each ROI. As expected, non-NK cell populations were also found to have changes in cell abundance due to inflammation, and we observed increases in FRCs, CD68+ macrophages, and CD4+ T cells (Fig. 7B; Supp. Fig. 6B). When assessing changes in NK cell subset abundance by CyCIF, we did not observe significant log2-fold differences in non-proliferating (Ki67–) NK progenitor abundance between non-inflamed and inflamed donors (Fig. 7B). Local inflammation resulted in a significant log2-fold decrease in the abundance of tissue resident (CD69+ or CD103+) stage 4 and stage 5 NK cells (Fig. 7B). On the other hand, the presence of local inflammation in tonsil led to significant increases in the abundance of non-tissue resident stage 4 and 5 NK cells (Fig. 7B). Additionally, all proliferating (Ki67+) NK cell subsets were more abundant in tonsils with local inflammation including NK progenitors and stage 4 NK cells (Fig. 7B). To validate these observations, we quantified the frequency of NK cell subsets in CD45+Lin– live cells from 6 non-inflamed donors and 9 inflamed tonsil donors by flow cytometry (Fig. 7C). NK cell subsets made up a larger proportion (>3%) of CD45+Lin–Live cells in donors with inflammation, leading to an almost 2-fold increase in total NK cell frequencies (Fig. 7C). When looking for the developmental subsets that were most contributing to this increase, we specifically found that the frequency of stage 4B and 5 NK cells were significantly elevated in donors with local inflammation (Fig. 7C, Supp. Fig. 6D).

**Fig. 7.**
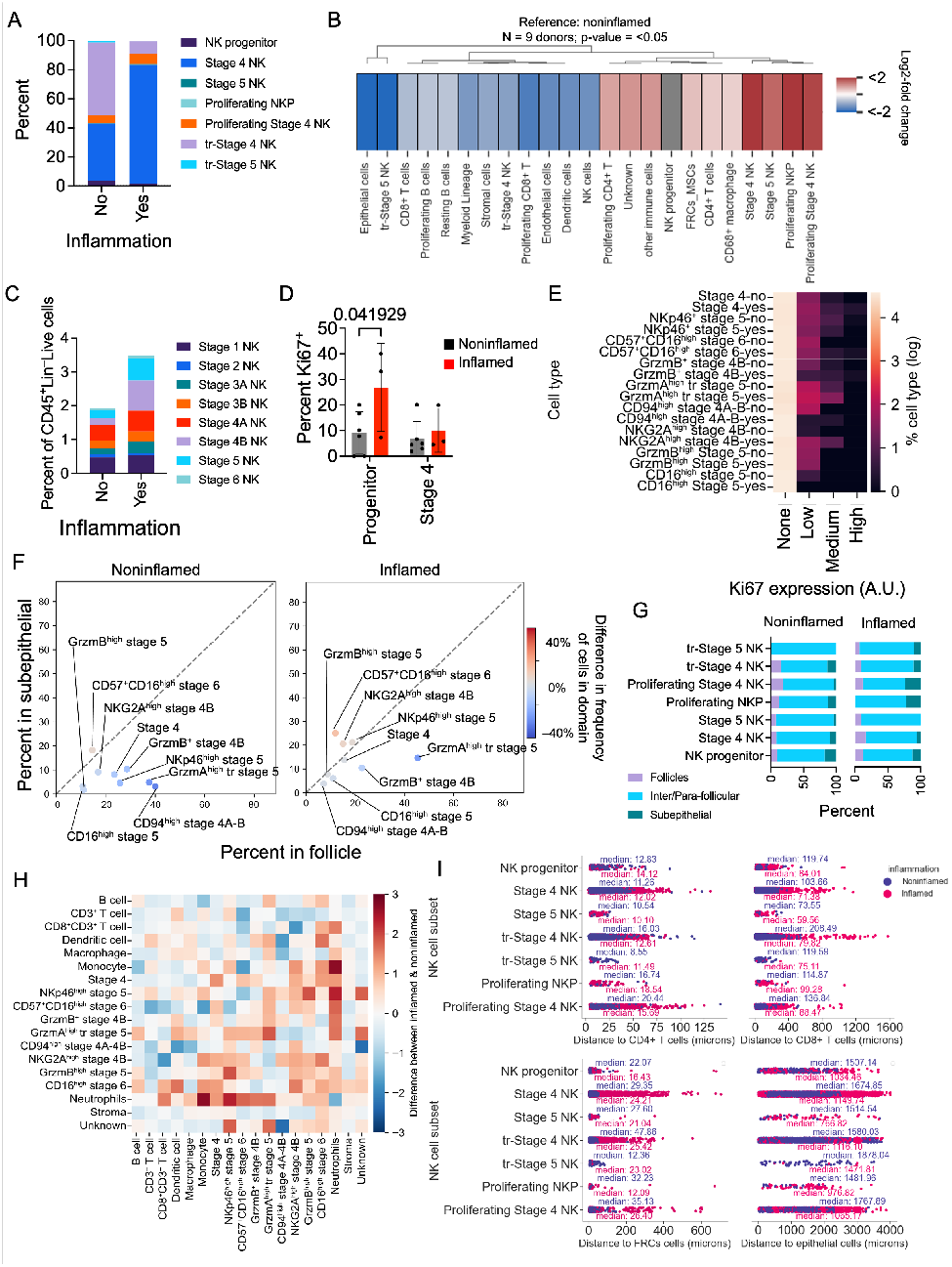
Inflammation affects NK cell subset frequency, localization, and interactions in tonsillar tissue. A) Frequency of NK cell subsets in inflamed (n=3 donors; 3 ROIs) and noninflamed (n=6; 6 ROIs) tonsil tissues calculated using Cy-CIF. B) Log2-fold difference in the abundance of tonsil NK cell subsets and other identified cell populations between inflamed donors (n=3) and non-inflamed donors (n=6). Log2-fold change was calculated relative to non-inflamed donors. Non-significant changes are shown in grey. C) Percent of NK cell subsets from 6 non-inflamed and 9 inflamed tonsil donors quantified from flow cytometric analysis. Tonsil mononuclear cells were isolated and stained with the antibody panel described in Supp. Table 4. D) The percent of Ki67+ NK cells identified by CyCIF from 6 non-inflamed donors (6 ROIs) and 3 inflamed donors (3 ROIs). p-values were calculated by one-way ANOVA with multiple comparisons. E) Comparison of the average percent of Ki67 positive NK cells from 2 inflamed and 3 non-inflamed donors using IMC data. Relative expression of Ki67 on Ki67+ NK cells were binned by log normalized expression intensity [bins=none, low, medium, and high]. F) Comparison of the average spatial localization of tonsil NK cells between 2 inflamed and 3 noninflamed tonsil donors. X-axis, percent of cells in follicle; Y-axis, percent of cells within the subepithelial domain; quantified from IMC derived single cell data. Color of data point represents the difference in the frequency of cells found in the same domain between inflamed and non-inflamed donors. G) Comparison of the frequency of NK cell subsets in each tonsil domain between 2 inflamed and 4 noninflamed donors calculated from CyCIF data (6 ROIs). Quantification of frequency of NK cell subsets in each domain was performed as described for Fig. 2B. H) Heatmap depicting the difference in cell-cell interactions between inflamed (n=2) and noninflamed (n=3) donors quantified from IMC derived single cell data. Increased frequency of a particular cell-cell interaction due to inflammation is represented by red and decreased cell-cell interactions are assigned blue. I) Median NK cell subset distance to CD4+ T cells, CD8+ T cells, FRCs, and epithelial cells quantified from CyCIF data of 3 inflamed and 6 non-inflamed donors (9 ROIs).

To further determine whether inflammation contributed to differential proliferation between NK cell subsets, we analyzed relative Ki67 expression in NK cell populations using CyCIF (Fig. 7D) and IMC data (Fig. 7E). The percentage of Ki67+ cells was significantly increased in the NK cell progenitor population with a trend towards an increase in the stage 4 NK cells (Fig. 7D). Nevertheless, the log2-fold change in Ki67+ stage 4 and 5 NK cell abundance in inflamed donors relative to non-inflamed donors suggests that these differences are biologically relevant (Fig. 7B). Additionally, analysis of Ki67 expression from our IMC data was used to define differences in proliferation between phenotypically distinct stage 4 and 5 populations. This analysis showed that the presence of inflammation was accompanied by increased proliferation of NKG2A^high^ stage 4B and GrzmA^high^ stage 5 NK cells (Fig. 7E). Interestingly, CD94^high^ stage 4A-B NK cells were more proliferative in settings without inflammation (Fig. 7E). Overall, the presence of local inflammation induces an increase in proliferating NK cell progenitors and thereby likely provide a larger pool of maturing NK cells contributing to the overall increase in NK cells observed.

To define whether the localization of NK cell subsets changed under inflammatory settings, we compared the spatial distribution of NK cell subsets in inflamed and non-inflamed samples (Fig. 7F, G). Spatial analysis of IMC data revealed that stage 4 and 5 NK cells are more abundant within the inter/para-follicular domains in the presence of local inflammation, which is marked by decreased frequency of NK cells in and near the subepithelial (y-axis) and follicle domains (x-axis; Fig. 7F). Using CyCIF, we quantified the percent and absolute count of NK cell subsets in each tonsil domain and compared donors with (n=3 donors) or without (n=6 donors) local inflammation (Fig. 7G; Supp. Fig. 6C). We found that in the presence of local inflammation there was a reduction in the frequency of non-tissue resident NK cells within the subepithelial and follicle domains (Fig. 7G). All NK cell subsets detected by CyCIF, including non-tissue resident, tissue resident, and proliferating populations, were most frequently detected within the interfollicular and parafollicular in the presence of inflammation (Fig. 7G). Local inflammation was also associated with an increase in proliferating NK cell subsets in the subepithelial domain (Fig. 7G). Together, this shows that local inflammation leads to increased abundance of NK cells in interfollicular and parafollicular domains.

Finally, we sought to identify cell-cell contacts that were altered by local inflammation. We first performed analysis of direct cell contacts and measured the difference in interactions between inflamed vs. non-inflamed conditions using our IMC data. This analysis identified interactions that were significantly increased when comparing inflamed to non-inflamed conditions (Fig. 7H). In settings of local inflammation there was no major difference in NK cell interactions with T cells except for CD16^high^ stage 5 NK cells, which had increased interactions with CD8+ T cells (Fig. 7H). Most notably, we observed increases in interactions between NK cell subsets and neutrophils (Fig. 7H). We also observed that stage 4B and 5 NK cell clusters had increased interactions with dendritic cells and monocytes and to a more varying degree with macrophages (Fig. 7H).

Using CyCIF, we found differences in cell-cell interactions due to inflammation by quantifying the shortest median distance of NK cells to each identified population as described in Fig. 3. We found that with local inflammation, tissue resident and non-tissue resident stage 4 and 5 were in closer proximity to CD8+ T cells which is indicative of increased interactions (Fig. 7I). No major changes were noted in CD4+ T and NK cell or FRC and NK cell interactions in the presence of local inflammation based on shortest median distance (Fig. 7I). Moreover, NK cells appeared to move towards epithelial cells, dendritic cells, and other myeloid cells in tonsils with local inflammation (Fig. 7I; Supp. Fig. 6E). Tissue resident stage 4 NK cells were the only subset that were localized closer to CD68+ macrophages by >30 µm in settings of local inflammation (Supp. Fig. 6E). In the presence of local inflammation, NK cells did not engage in novel cell-cell interactions rather we noted differences in the likelihood of cell-cell interactions.

Overall, the presence of local inflammation in tonsil increases the proliferation and abundance of NK cell subsets in tonsil. Specifically, we see that NK cells make up a larger proportion of CD45+ lineage negative (CD3, CD14, CD19) cells in inflamed tonsils, with non-tissue resident stages 4 and 5 NK cells contributing the most significant increases in abundance. The localization of NK cells is also seen to become more restricted to the parafollicular and interfollicular domain.

## Discussion

While the phenotypic and transcriptional trajectory of human NK cell development has been described, the spatial niches that support human NK cell development have not been well studied (2). Accurate identification of NK cell developmental subsets requires the quantification of multiple surface receptors, limiting the use of conventional fluorescence-based imaging techniques (1). Using IMC and CyCIF, our study provides comprehensive spatially resolved single cell analysis of NK cell developmental subsets in pediatric tonsils and serves as a resource for future studies, particularly when paired with recently published single cell atlases of human tonsil immune and non-immune subsets (47–49). IMC and CyCIF panels enabled us to identify and phenotype NK cell developmental subsets in pediatric tonsil. Broadly, we found that NK cell localization and cell-cell interactions are dependent on tissue residency and NK cell developmental stage. We sought to define the tissue niches that support NK cell differentiation, particularly the sites of earliest stages of NK cell development. NK cell maturation in tonsil is thought to begin with a circulating precursor that enters tissue from peripheral blood (1, 4, 34). It is conceivable that an SLT tissue-seeding precursor has the same, or similar, identity as bone marrow-derived tissue-seeding precursors that enter the thymus from peripheral blood and give rise to mature thymocytes and other immune lineages, including mature NK cells, in postnatal T cell homeostasis (50, 51). Although the signals that mediate homing of circulating CD34+ NK cell progenitors to SLT have not been defined, their expression of chemokine receptors CXCR4 and CCR7, which can interact with CXCL12 and CCL21 respectively, suggests that they enter via HEV or blood vessels expressing these chemokine ligands (30, 46, 52). This observation is additionally supported in our data showing proximity of precursors to blood vessels within the interfollicular domain. How, or if, precursors are targeted to thymus or SLT, and similarly whether lineage fate is encoded before their recruitment or occurs solely from stochastic signals in the environment, remains to be determined for human NK and T cell precursors (53).

NK cell progenitors have been observed by single marker IHC staining to localize to the interfollicular and parafollicular domains within tonsil (9). We identified CD34+CD45+CD122+ progenitors expressing CD49d, the integrin alpha chain associated with integrin β7, in the interfollicular and parafollicular domain adjacent to stromal cells, including FRCs. While we did not explicitly test CD45RA expression, the expression of CD45 and CD49d suggests that these cells are analogous to previously described CD34+CD45RA+integrin β7bright precursors. Direct interaction with developmentally supportive stromal cells improves the efficiency of NK cell maturation from CD34+ progenitors in vitro, suggesting that interactions between NK cell subsets and stromal cells or FRCs similarly supports particularly early stages of NK cell development in vivo (21, 54–56). Our shortest distance analysis validates this hypothesis, as it suggests that early (stage 1-2) NK progenitors receive cytokines through direct cell-cell interactions with FRCs.

While stage 1 and 2 CD34+ NK cell progenitors are rare in tonsil, our flow cytometry and imaging data show they are the most proliferative NK cell developmental subset, reflective of their proximity to cytokine producing FRCs. Interactions with FRCs presenting membrane-bound cytokines including IL-7, SCF, Flt3L, and IL-15, could thus lead to proliferation and maturation of NK cell progenitors (25, 40–42, 57). The expression of high affinity IL-2 receptor on NK cell progenitors in tonsil suggests that they also can respond to IL-2 from T cells, like mature NK cell populations (9). While NK cell progenitors likely receive proliferation signals from FRCs, proliferating mature NK cell populations are less likely to be in direct contact with FRCs. As such, we propose that the trafficking and subsequent proliferation of rare tissue-seeding progenitors can provide a larger pool of stage 3 NK cell precursors that can continue through maturation to become mature ILC or NK subsets following additional differentiation and maturation signals (5).

The localization of NK cell progenitors to the parafollicular and interfollicular domains, where FRCs are found, may also be directed in part by CXCL12 and CCL21, as stage 1/2 NK progenitors expressed CXCR4 and CCR7 and were in proximity to CXCL12+ and CCL21+ cells. While CCR7 was poorly detected by flow cytometry on most NK cell subsets, its gene expression and detection on stage 2 NK cells suggests that we detected this NK progenitor by CyCIF in tissue, and previous studies have reported responsiveness of CD56+CD16– cells to CCR7 ligands (52, 58). CXCR5 expression on NK cell progenitors could also direct them towards the follicle, but more likely is important for trafficking into tonsils via CXCL13+ HEVs (15). As NK cells progress though maturation from progenitor to stages 4 and 5, they move away from the interfollicular domain to the parafollicular T cell rich zone. This observation confirms previous observations of NK cells in the parafollicular microdomain where they can respond to IL-2 from T cells (12). Most proliferating stage 4 and 5 NK cells are not in proximity to IL-7, IL-15, or Flt3L expressing cells, suggesting that other factors, including IL-2, can promote mature NK cell proliferation in situ (12). In addition, the increased frequency of interactions between mature NK cells and CD4+ T cells underscores the role of NK cells in assisting in the coordination of adaptive immunity (59–61). Finally, while these data identify certain correlations between chemokine receptors and ligands, it should be noted that trafficking in tissue is complex and directed by chemokine signaling, cell adhesion, and physical properties of tissue.

Although we predicted that tissue resident NK cell populations would have unique localization compared to non-tissue resident NK cells, we observed similar abundance of tissue resident and non-tissue resident NK cells in each domain. Still, within the interfollicular and parafollicular domains tissue resident and non-tissue resident NK cell subsets occupy unique spaces based on their distinct cell-cell interactions and proximity to chemokine ligands including CXCL12, CCL19, and CCL21. Tissue resident NK cells have more interactions with macrophage and other myeloid lineage cells than non-tissue resident NK cells, suggesting tissue resident populations are more likely to be generating or receiving signals from myeloid cells. Non-tissue resident NK cells, although they can interact with myeloid cells, are observed to preferentially interact with T cells, suggesting they are modulating T cell responses and elimination (62). NK cells can influence T cell responses in antiviral and inflammatory settings in part preventing autoimmunity by eliminating activated CD4+ and CD8+ T cells via perforin-dependent killing, and TRAIL and Fas pathways (60, 63–65). Moreover, we show that these interactions become more prevalent in the presence of local inflammation, further indicating their importance in coordinating innate and adaptive immune responses.

The anatomical position of tonsils makes them the first line defense against inhaled and ingested pathogens and prone to inflammation. Local inflammation in tonsils can become chronic and is linked to changes in lymphocyte populations in response to pro-inflammatory cytokines (66, 67). Specifically, B cells and ILCs are dysregulated in pediatric patients with chronic local inflammation, suggesting that NK cells could also be affected (66). Increased localization of NK cell subsets to the interfollicular and parafollicular domain in the presence of local inflammation could be attributed to multiple factors. First, increased proliferation in mature and progenitor NK cells in these domains could contribute to elevated NK cell numbers and frequency. Second, increased recruitment of NK cell subsets to the interfollicular and parafollicular domains from other tonsil domains could cause increased abundance in the parafollicular domain. Third, based on the observed increased frequency of non-tissue resident population in tonsils with local inflammation, there is likely increased infiltration of circulating subsets into interfollicular and parafollicular domains where HEVs are more present. In addition, it remains to be determined how NK cell localization, interactions, and development, in SLT, are shaped by specific disease settings, including viral infection and malignancy.

In summary, this resource is the first comprehensive analysis of human NK cell developmental subsets in situ and provides insights into the function, developmental niche, and trafficking of NK cell subsets. Based on our observations, circulating progenitor cells enter the tonsil via HEVs lined with CXCL12+ and CCL21+ endothelial cells in the inter-follicular domain. Once there, NK cell progenitors localize to FRCs which present cytokines that induce proliferation and maturation. As progenitors mature into tissue resident and non-tissue resident stage 4 and 5 NK cells, they migrate to the parafollicular and subepithelial domains, where they can engage in interactions with T cells and myeloid cells. In the presence of local inflammation, NK cells are more proliferative and more abundant within the parafollicular domain, where non-tissue resident NK cells interact with T cells and tissue resident NK cells interact with both T cells and myeloid cells. Together, our study defines a road map for human NK cell development at steady state and under inflammatory conditions.

## Methods

### Tissue acquisition and processing

Matched peripheral blood and tonsillar tissue were collected from pediatric patients undergoing routine tonsillectomies at Columbia University Irving Medical Center. All tissue and blood samples were collected in accordance with the Declaration of Helsinki, with written and informed consent from all participants under the guidance of the Institutional Review Board of Columbia University. Post-operative pathology reports were used to determine whether local inflammation was present. Tonsillar tissue samples were split in two sections to make single cell suspensions and formaldehyde fixed paraffin embedded (FFPE) tissue slides. To produce single cell suspensions from tonsil, specimens were placed into a petri dish with sterile PBS and manually dissociated by mincing the tissue into a cell suspension which was then filtered through a 40 µm cell strainer. Samples were then centrifugated at 1200 rpm for 7 minutes and resuspended in PBS followed by another round of centrifugation. Tonsillar cell suspensions were resuspended in heat inactivated FBS with 10% DMSO to cryopreserve for flow cytometric analysis. The remaining tonsil specimens were fixed in 10% paraformaldehyde for 24-48 hours then placed in 70% ethanol. Fixed specimens were given to the Herbert Irving Comprehensive Cancer Center Histology Service and Tumor Banking core for paraffin embedding, tissue sectioning, and slide preparation. 3 µm sections were prepared for imaging mass cytometry and high-resolution microscopy, and 10 µm sections for spatial transcriptomics.

Blood from pediatric tonsil donors were collected by venipuncture on the day of their tonsillectomy. Whole blood was diluted with equal parts of PBS and layered on FicollPaque density gradient followed by centrifugation at 2,400 rpm for 20 mins with no brake and slow acceleration. The lymphocyte layer was collected from gradient interface and washed with equal parts of sterile PBS then centrifuged at 1,200 rpm for 5 mins. Collected cells were resuspended in HI FBS with 10% DMSO at 1-5 x 10^6^ cells/mL and cryopreserved for flow cytometric analysis.

### Spatial transcriptomics

FFPE tissue was prepared into 10 µm sections and sent to Nanostring for GeoMx RNA assay data acquisition on a GeoMx Digital Spatial Analyzer (DSP). 35 domains of interest were chosen based on tonsillar microarchitecture using tissue sections from 6 pediatric donors (4 noninflamed and 2 inflamed). Spatial transcriptomic data was normalized using Q3 normalization after BioQC and filtering using GeoMx DSP Analysis Suite (version 2.5.1.145). Heatmaps were produced using ClustVis (68) and Heatmapper (69).

### Flow cytometry and analysis

Cryopreserved tonsillar cell suspensions and peripheral blood mononuclear cells were thawed and resuspended in sterile PBS with 10% FBS. Before immunostaining, antibodies specific to extracellular proteins were diluted in PBS with 10% FBS (Supplemental tables 3-4). Cells were stained for extracellular markers for 30 mins at room temperature (RT) then washed. If required, cells were then fixed and permeabilized using eBioscience FoxP3 buffer (catalog no. 00-5523-00; ThermoFisher) then immunostained for intracellular markers for 30 mins at RT. A Novocyte Penteon Cell Analyzer was used to acquire data, which was then exported to FlowJo (BD Biosciences) or OMIQ (Dotmatics) for downstream analysis. For Ki67 expression and subset frequency calculations, NK cell developmental subsets (stage 1-6) were identified using the gating strategy described in Supp. Fig. 7A and B. For defining chemokine receptor expression of NK cell developmental subsets, we identified NK cell developmental subsets using single-cell analysis in OMIQ. First, data were arcsinh transformed and live single CD45+lin— cells were gated. FlowAI (70) was applied to remove low quality cells and FlowSOM (71, 72) based on all the markers of the backbone panel was applied to remove any residual lineage positive dead cells and innate lymphoid cells (CD127+NKp44+, CD127+CD103+, and CD127+CD294+). On the remaining cells a similar gating strategy was applied as described in Supp. Fig. 7A and Chemokine receptor positive gating on NK cell subsets was determined by fluorescence minus one control. For flow cytometric analysis of human stromal cells from bone marrow and tonsil, data were arcsinh transformed then cells were gated to remove lineage positive (CD14+, CD19+, CD45+, CD3+) and dead cells before performing UMAP dimensional reduction analysis using all markers.

### IMC and CyCIF microscopy

FFPE tonsil sections were stored at 4C and brought to room temperature prior to dewaxing with three 10-minute Xylene (catalog no. X5–1; Fisher Scientific) washes. Next, tissue was rehydrated using serial washes in 100% and 95% ethanol (catalog no. BP2818100; Fisher Scientific) for 10-minutes, then in 70% and 35% ethanol for 5 minutes. Excess ethanol was washed out with water. Heat-activated antigen retrieval was carried out on rehydrated tissue sections by incubating slides in prewarmed Agilent Dako pH6 antigen retrieval solution (catalog no. S2367; Agilent) for 20-minutes in a vegetable steamer (Black &Decker), cooled to room temperature, and washed with PBS. For CyCIF, tissue sections were incubated with 1 mg/mL borohydride in deionized water for 10 minutes to decrease background fluorescence then washed three times with PBS. Samples were permeabilized with 0.2% Triton X-100 in PBS for 30 minutes and washed with PBS three times. After permeabilization, tissue samples were incubated with ThermoFisher Super Blocking Buffer (catalog no. 37580; Thermo Fisher) for 1 hour. Antibodies for immunostaining were diluted in antibody diluent buffer [PBS/10% Bovine Serum Albumin/0.01% Triton X-100], dilutions for each antibody can be found in Supplemental tables 1 (IMC) and 2 (CyCIF). Unconjugated primary antibodies for IMC were first conjugated to polymer and labelled with a unique metal isotope using Maxpar Antibody Labelling Kit (catalog no. 201300; Standard BioTools [Fluidigm]). Immunostaining of tissue sections with unconjugated antibodies for high-resolution immunofluorescence microscopy and with metal-conjugated antibodies for IMC was done overnight at 4C in a humidity chamber.

For IMC, tissue sections were first stained with NK cell specific antibodies overnight then washed with antibody diluent buffer and stained for the remaining markers overnight. After immunostaining, tissue sections were washed five times with antibody diluent buffer and twice with PBS. Cell-ID Intercalator-Ir 125 µmolar (catalog no. 201192A; Standard BioTools) was diluted 1:400 in PBS then added to tissue samples for 45 minutes at room temperature. Tissue was subsequently washed four times with DI water and then allowed to dry at room temperature. IMC data were acquired using a Hyperion Imaging System.

For immunofluorescence microscopy, unbound antibodies were washed with antibody diluent buffer four times after immunostaining. Tissue sections were incubated with secondary antibodies at a dilution of 1:200 for 1 hour at room temperature followed by four washes with antibody diluent buffer and two washes with PBS. Washing buffer was removed from tissue prior to adding ProLong Glass Antifade Mountant with NucBlue Stain (catalog no. P36985; ThermoFisher) and a #1.5 coverslip (catalog no. 72204-03; Electron Microscopy Services). Samples were cured overnight at room temperature prior to imaging with a Zeiss CellDiscoverer 7 using a 20x/0.7 NA objective and an optical magnification of 0.5x. For CyCIF microscopy, tissue was treated with Vector TrueVIEW Autofluorescence Quenching Kit (catalog no. SP-8400; Vector Labs) prior to mounting. After washing excess TrueView reagents, tissue sections were mounted using VectaShield Antifade Mounting Medium with DAPI (catalog no. H-1200; Vector Laboratories, Inc) and a #1.5 coverslip (catalog no. 72204-03; Electron Microscopy Services). domains of interest were imaged with a 20% overlap using a Zeiss CellDiscoverer 7 using a 20X/0.7 or 20X/0.95 NA objective and an optical magnification of 0.5X and coordinates were saved for subsequent imaging. This resulted in images with a micron per pixel unit of 0.454 microns/pixel. Upon completion of imaging, coverslips were gently removed by placing slides in a Coplin Jar with PBS. To strip antibodies, tissue slides were incubated in freshly prepared stripping buffer (20 mL 10% SDS/0.8 mL beta-mercaptoethanol/12.5 mL 0.5 M Tris-HCl (pH 6.8)/67.5 mL DI H2O) for 30 mins at 65C then washed with water for 15 mins. Slides were then blocked and re-stained with primary and secondary antibodies prior to re-imaging as described above.

### CyCIF and IMC data analysis

Illumination correction was applied using BaSiC darkfield and flat-field profiles (39, 73). Stitching of tiles and alignment of channels was done using Ashlar with a barrel correction [value used: 7.943e-09]. Background subtraction (Backsub) module was applied to reduce autofluorescence in all channels except for collagen, CD10, CD3, and IL-15. These channels were omitted from background subtraction as the signal to noise ratio was not high enough. Nuclei-based cell segmentation was performed based on DAPI nuclear staining from first cycle using HALO AI (Indica Labs) or Mesmer (39, 74). CD20, CD127, CD45, CD10, and CD3 were used to define cell membrane boundaries for Mesmer membrane-based cell segmentation. Single-cell data was extracted using McQuant. All packages, except Ashlar, were applied using the McMicro pipeline (39, 73). Subsequent single-cell analysis was performed with SCIMAP (https://scimap.xyz/). For cell phenotyping, CyCIF fluorescence intensity data was first scaled based on manual gating which defined positive and negative intensity thresholds for each marker. The scaled data was then used to define populations based on a phenotype workflow which provided the marker expression associated with each population. NK cells were identified using the strategy in Supp. Fig. 3A after excluding lineage positive cells (Lineage=CD3, CD20 CD31, smooth muscle actin (SMA), gp38, pan-keratin, collagen type 1). For quantifying cell abundance using SCIMAP, tonsil microdomains were manually annotated based on the marker intensity of CD3, CD20, CD10, collagen type 1, smooth muscle actin, pan keratin, and gp38. HALO AI network was trained by manually defining representative domains and considered all marker intensities to identify domains and quantify CXCL12, CXCL13, CCL19, CCL20 and CCL21 expression. SCIMAP was also used for measuring log10-fold difference in cell-type abundance, spatial co-occurrence analysis and calculating distance between cell types. Spatial co-occurrence analysis shown in Fig. 3B was calculated using the radius method to measure all cells within a 50-pixel (22.7 µm) radius from the center of each cell to get the probability and significance of cells colocalizing. Average shortest distance function was used to measure the distance between cell types and takes the average of the shortest distance [microns] between each cell and cell type. Microscopy data was visualized using Fiji (75, 76), QuPath (77), and Indica Labs HALO AI software. Prism 9 (GraphPad) and Seaborn [Python version 3.9] was used to plot and statistically analyze CyCIF data (78, 79).

Imaging mass cytometry data was preprocessed starting with MCD files and then converted to OME-TIFF format. Cell segmentation (Supp. Fig. 1B) was performed with a pretrained pixel classifier trained to distinguish nuclei (marked by DNA and or Histone H3 signal), cytoplasm (surrounding nuclei and overlapping cytoplasmic channel signal), and background (pixels with low intensity channels) using ilastik (version 1.3.3) (80). Class probabilities were fed to Deep-Cell (version 0.12.1) for segmentation of nuclear and cytoplasmatic compartments (81). Signal was quantified for each channel in each segmented cell and morphological features such as area, and eccentricity were also calculated for the cell masks. Values were centered, scaled and data integrated using harmony based on initial PCA dimensions (82). A nearest neighbor graph using the harmony integration was built, and a Uniform Manifold Approximation and Projection (UMAP) (version 0.5.3) was constructed for visualization. Finally, we clustered cells with the Leiden algorithm (leidenalg package, version 0.8.10) with resolution 3.0 (83). The Leiden clusters were labeled with a cell type label based on the abundance of signal from the channels (Supp. Fig. 2). Leiden clustering with resolution 0.5 was also applied to the NK cell compartment to identify unique subsets. All steps were performed with Scanpy (version 1.9.1) (84). To provide a microanatomical context to the single-cell data, we used UTAG to derive areas of tonsil microanatomy in an unsupervised manner using graphs (85). We labeled these clusters into the following classes: follicle, follicle adjacent area, interfollicular/parafollicular area, subepithelial and blood/lymphatic vasculature. We sought to go beyond simple association of cells with microanatomical labels, but also performed instance segmentation of microanatomical domains by observing connected components of each domain type in each region of interest (ROI) and kept instances made up of at least 0.5% of cells in each ROI. This allowed us to identify specific instances of domains and pinpoint their centroid location which will be important in measuring distances of cells to specific domain instances and the enrichment in each microanatomical domain derived by UTAG. We then calculated overlap ratios of cells in microanatomical domains, but also measured the distance of every cell to every microanatomical domain centroid and border. Cell type interactions were determined based on physical proximity between cell types using a cell type label randomization strategy for background as previously described (86). Comparisons of marker expression were performed with Scanpy’s “rank_genes_groups” function using a T-test with adjustment for overestimation of variance. All comparisons of IMC data such as cell type abundance or marker expression were performed with a two-tailed Mann-Whitney U-test and adjusted for multiple testing with the Benjamini-Hochberg FDR method.

## ACKNOWLEDGEMENTS

The authors wish to thank research coordinators Evelyn Hernandez and Carlos Aguilar Breton for their assistance with the acquisition of tonsillar tissue from healthy donors. We would like to acknowledge the Columbia Stem Cell Initiative Flow Cytometry Core Facility at Columbia University Irving Medical Center, under the leadership of Michael Kissner, which was used to perform all flow cytometric analysis and cell sorting. We would also like to thank the Herbert Irving Comprehensive Cancer Center Histology Services and Tumor Banking core at Columbia University Irving Medical Center for assistance with tissue embedding and sectioning. This work was partially supported by the WorldQuant Initiative for Quantitative Prediction. We would like to thank the Englander Institute of Precision Medicine Mass Cytometry Core facility for providing labeling guidance and acquiring samples for IMC imaging. This work was in part supported by the NIH-NIAID R01AI137073 to EMM and F31AI164869 to EHS. A.F.R. was supported by NIH-NCI T32CA203702 grant until April 2022, and is supported by Angelini Ventures S.p.A. Rome, Italy since June 2022. B.I. was supported by NIH-NCI R37CA258829 and R01CA266446. B.I. is a consultant for or received honoraria from Volastra Therapeutics, Johnson &Johnson/Janssen, Novartis, Eisai, AstraZeneca and Merck, and has received research funding to Columbia University from Agenus, Alkermes, Arcus Biosciences, Checkmate Pharmaceuticals, Compugen, Immunocore, and Synthekine. A.G.F. is a consultant for ImmunoVec, Inc and Immunebridge, Inc.

