## Supplemental Figures for "Mapping human natural killer cell development in pediatric tonsil by imaging mass cytometry and high-resolution microscopy"

A

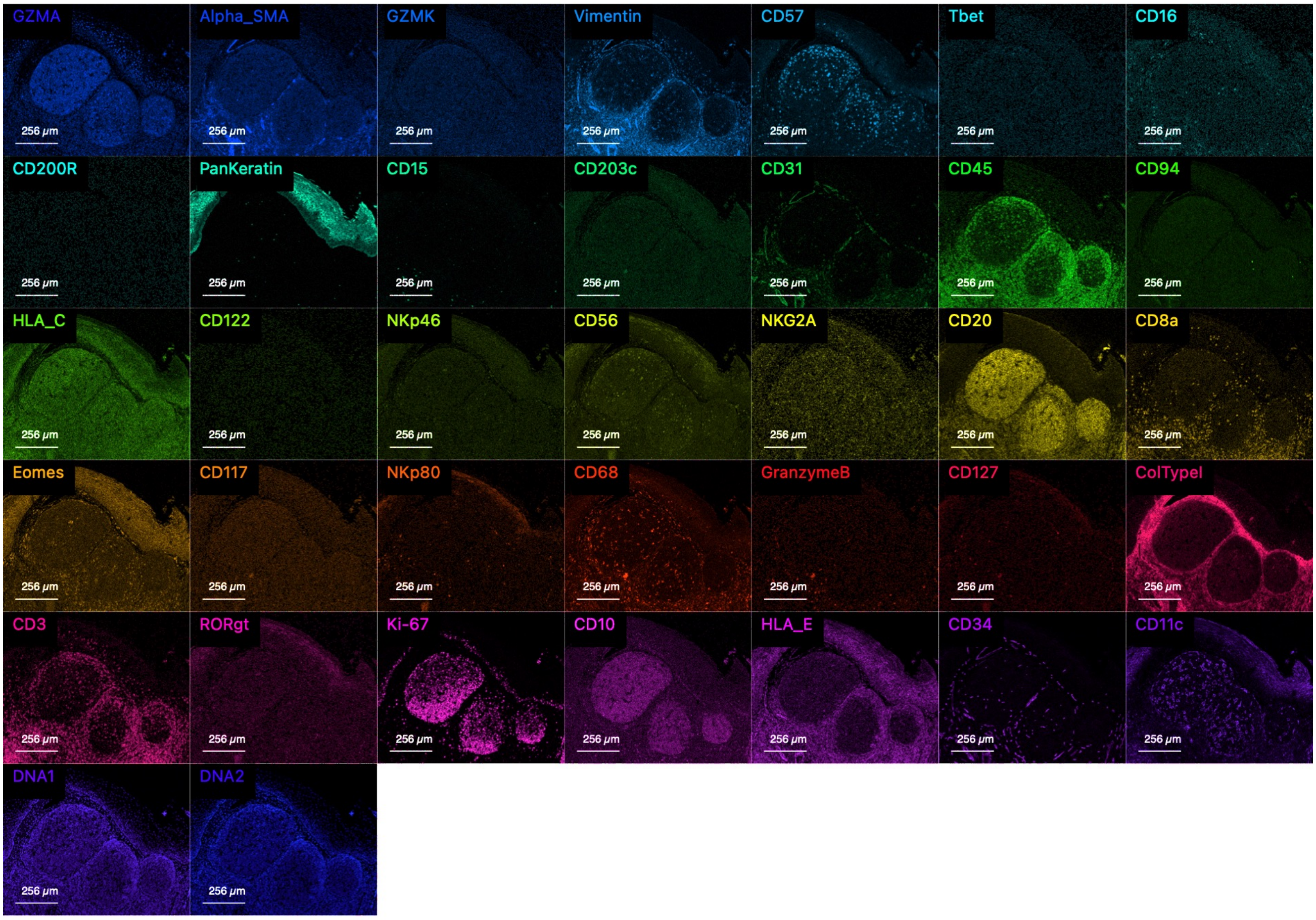

B

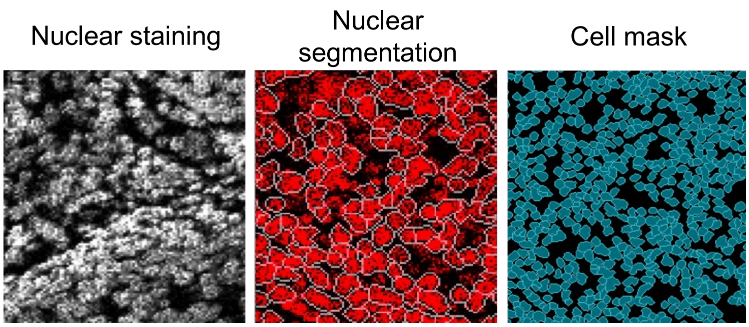

Supplemental Figure 1: A) Representative images of all markers captured by IMC for a single ROI. FFPE tonsil sections were immunostained with 35 metal-conjugated antibodies and 191/193-Ir DNA intercalator for IMC. B) Representative images of IMC nuclear staining, segmentation, and cell mask. Nuclear segmentation was performed based on 191/193-Ir DNA intercalator nuclear staining as described in the methods.

Supp. Figure 2

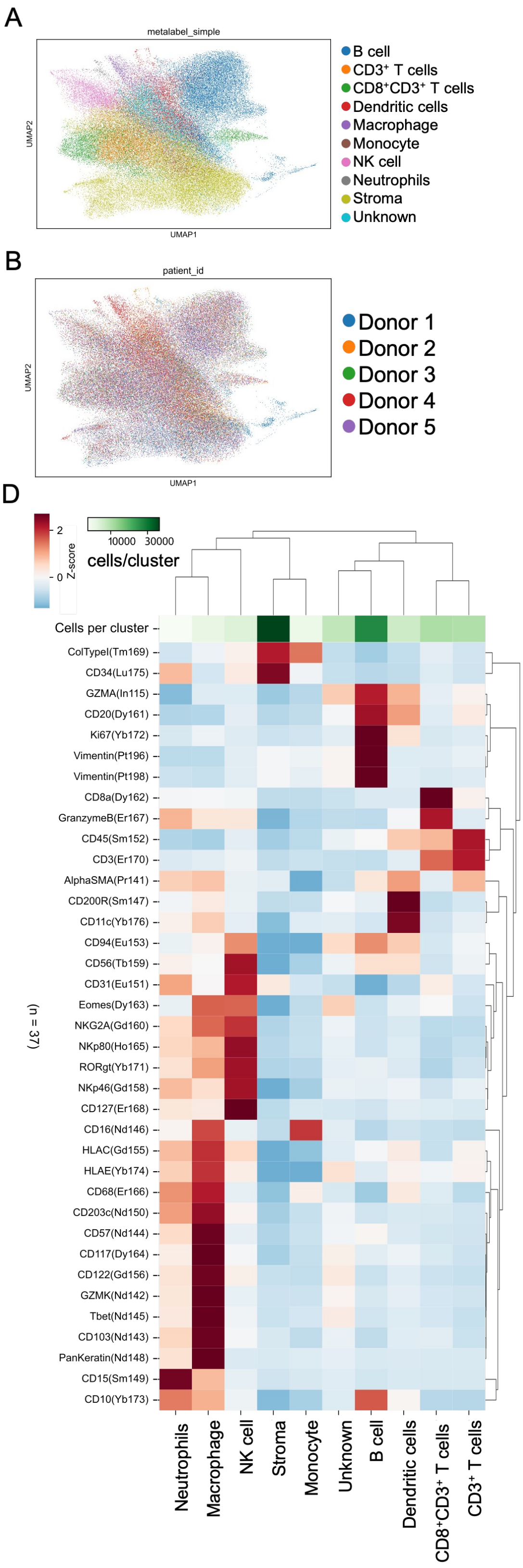

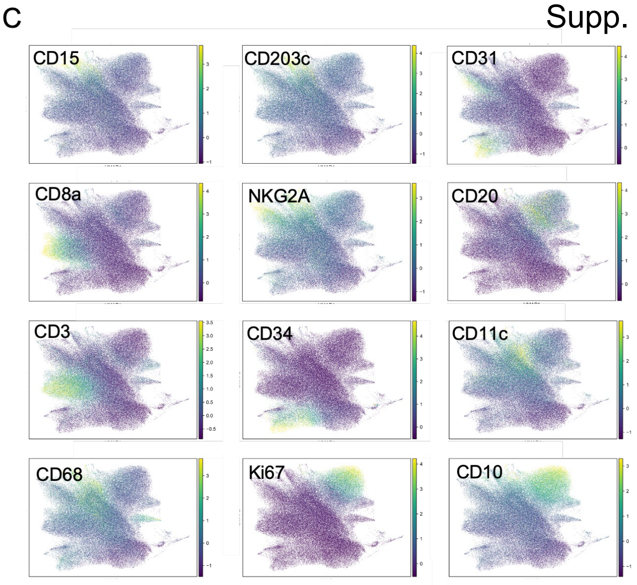

E

| Cell type | Markers |
|---|---|
| B cells | $CD45^+CD20^+CD10^{+/-}$ |
| CD8⁺CD3⁺ T cells | $CD45^+CD3^+CD8^+GZMB^+$ |
| CD3⁺ T cells | $CD45^+CD3^+CD8^-GZMB^-$ |
| Dendritic cells | $CD45^+CD11c^+CD3^-CD20^-$ $CD56^-$ |
| Monocytes | $CD45^+CD16^+CD3^-CD20^-CD56^-$ |
| Stroma | $CD45^-CD34^+CD31^+ColType1^+$ |
| NK cells | $CD45^+CD3^-CD56^+NKp46^+$ $NKG2A^+CD94^+$ |
| Macrophage | $CD45^+CD68^+CD3^-CD20^-CD56^-$ |
| Neutrophils | $CD45^+CD15^+CD3^-CD20^-CD56^-$ |

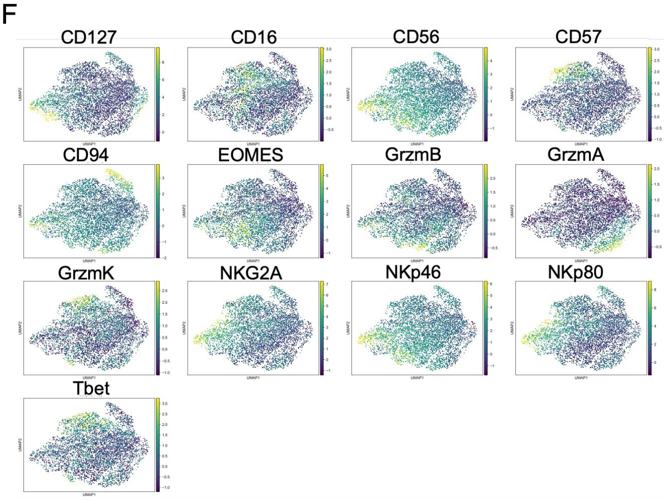

Supplemental Figure 2: A-C) UMAP projection of tonsillar cell populations identified by IMC and colored by (A) cell populations, (B) donor ID, and (C) marker expression intensity of those used for cell identification. D) Expression of 35 unique markers used to immunophenotype tonsillar cell populations from IMC data (n=5 donors). E) Strategy for identifying cell phenotypes of tonsil cell populations by their expression intensity of markers from the IMC panel. F) UMAP visualization of tonsil NK cell subsets identified from IMC data of 5 donors and colored by marker expression intensity.

Supp. Figure 3

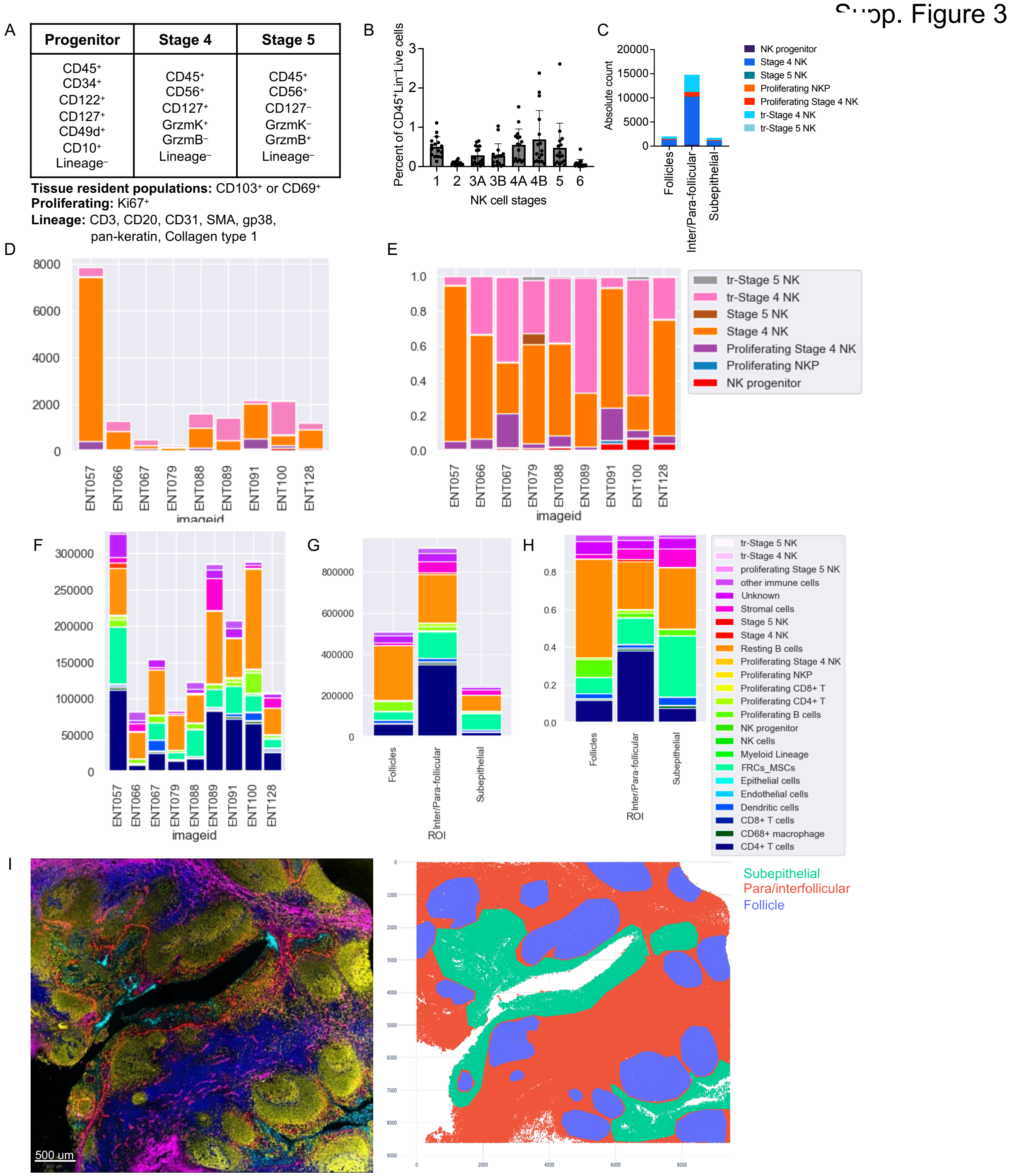

Supplemental Figure 3: A) Gating strategy used to identify NK cell populations by CyCIF. B) Percentage of NK cell developmental subsets that make up tonsil CD45⁺lineage⁻Live cells. Percent was calculated from flow cytometric analysis of 15 tonsil donors. C) Absolute count of NK cell subsets that were identified in each tonsil microdomain by CyCIF. Data was acquired from 9 donors and 9 ROIs. Domains were defined manually based on marker intensity and microarchitecture. D) Absolute count of NK cell subsets observed in donors used for CyCIF. E) Proportion of NK cell subsets that were identified in each donor by CyCIF (n=9 donors; 9 ROIs). F) Absolute count of all identified tonsil cell populations from CyCIF (n=9 donors; 9 ROIs 9=). G) Absolute count and (H) proportion of all cell populations in tonsil domains (n=9 donors). I) Representative CyCIF image of FFPE tonsil (left) and respective mask of doamin annotations (right) including subepithelial (green), follicle (blue), and interfollicular/parafollicular (orange) domains. CyCIF image shows markers used for defining domains including gp38 (red), CD3 (blue), CD20 (yellow), pan-keratin (cyan), and collagen type 1 (magenta).

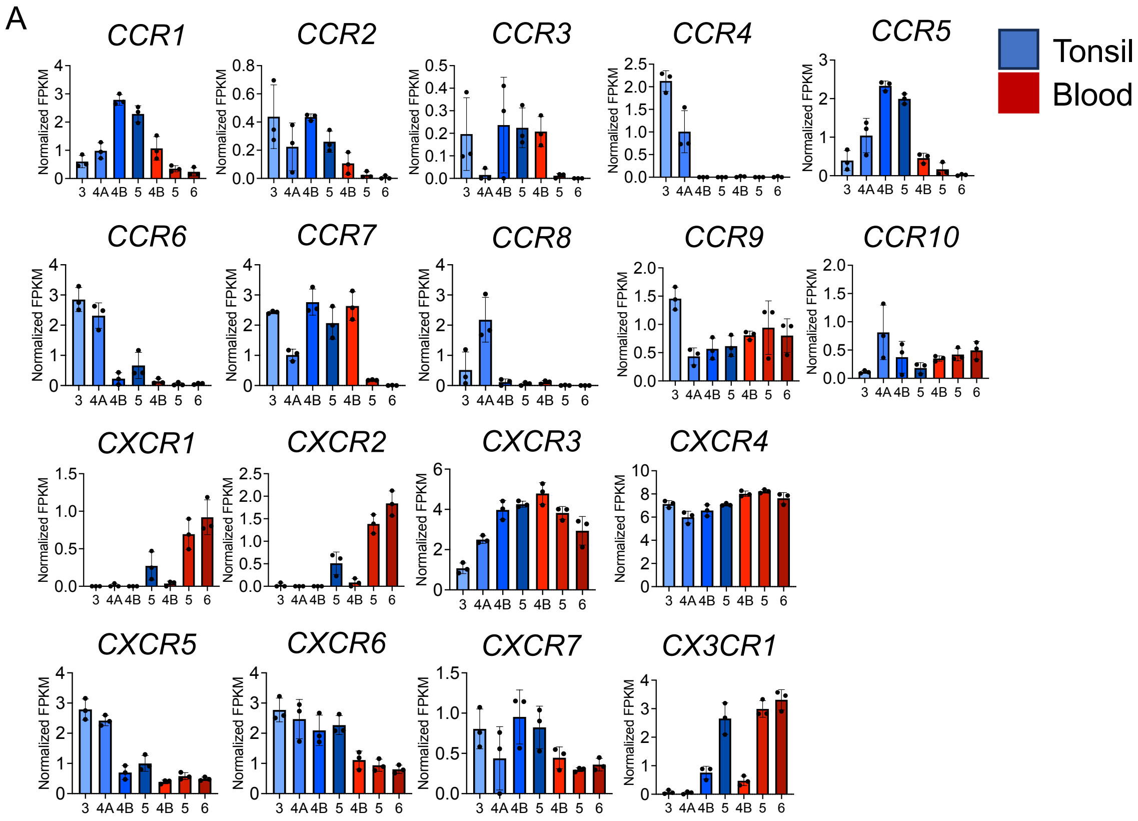

Supplemental Figure 4: A) Chemokine receptor gene expression (normalized FPKM) of sorted NK cell subsets from peripheral blood and tonsil. NK cell subsets were sorted by FACS and pooled into 3 technical replicates from 12 tonsil and peripheral blood donors for bulk RNA sequencing (Hegewisch-Solloa et al., 2021).

Supp. Figure 5

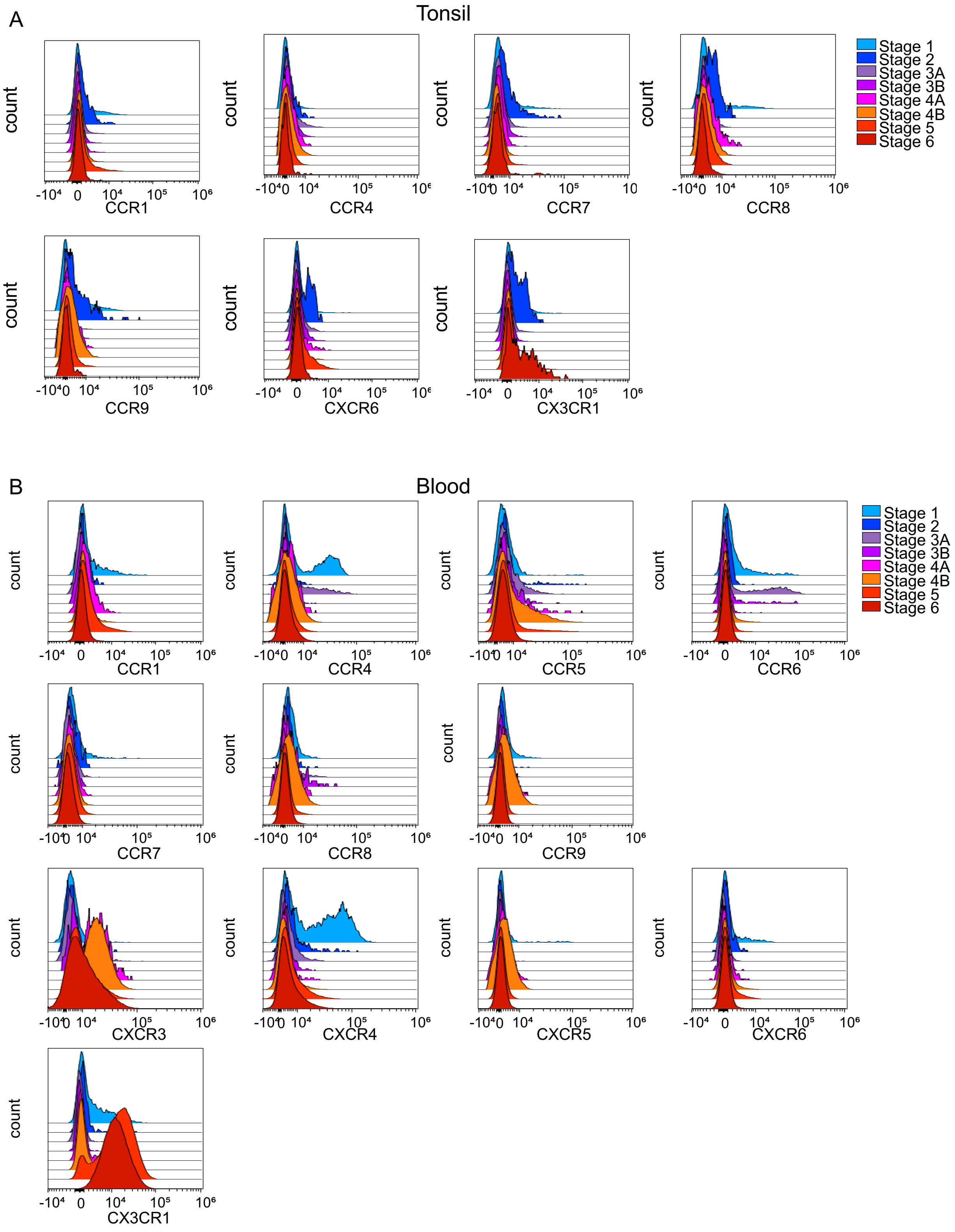

Supplemental Figure 5: A) Tonsil NK cell subset chemokine receptor expression measured by flow cytometry. Each histogram represents 6-7 donors. B) Peripheral blood NK cell subset chemokine receptor expression measured by flow cytometry. Each histogram represents 6-7 donors. Paired peripheral blood and tonsil donors were used for flow cytometric analysis of NK cell chemokine receptor expression.

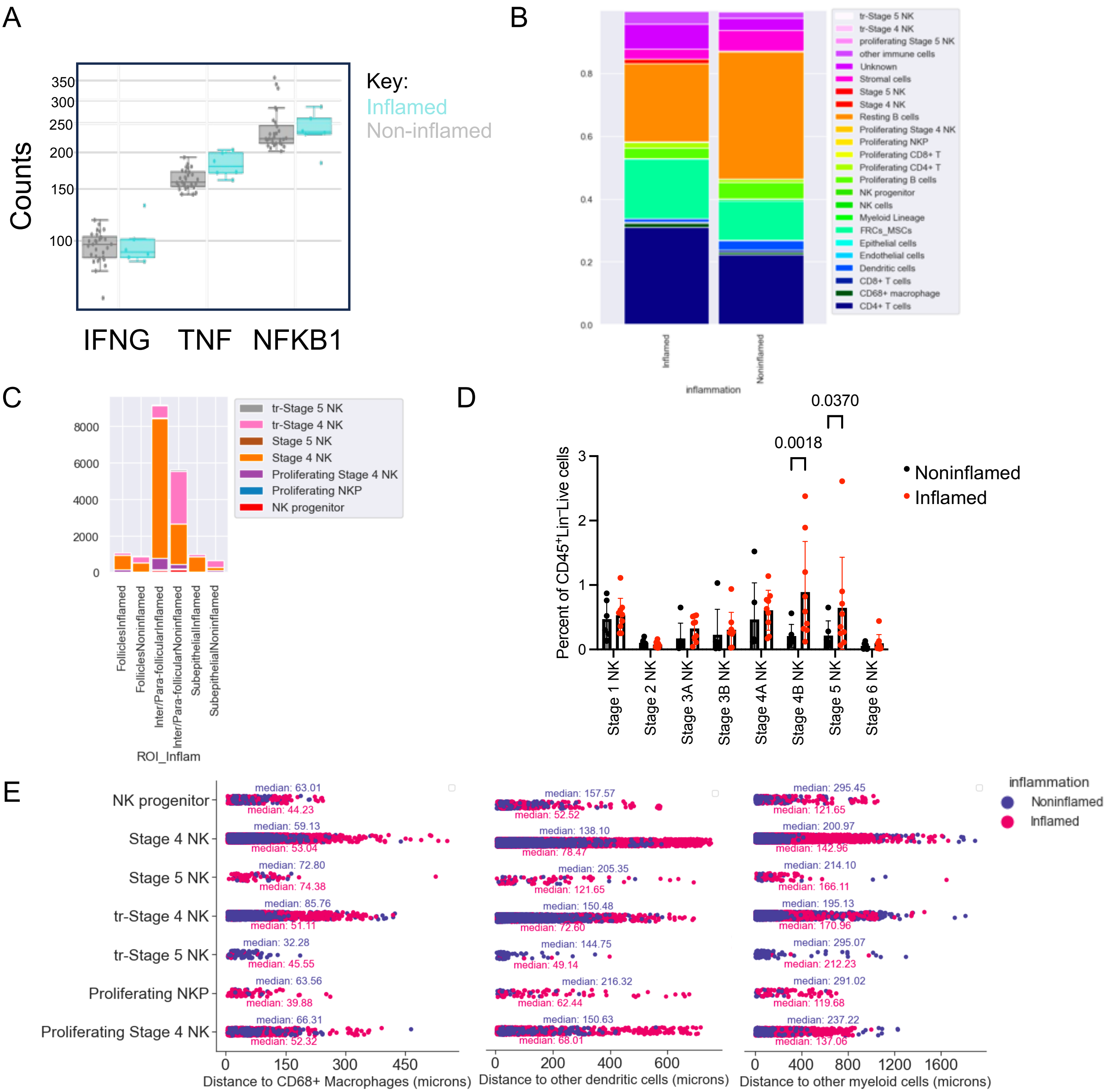

Supplemental Figure 6: A) *IFNG, TNF,* and *NFKB1* gene expression from Nanostring GeoMx of 32 ROIs from 4 noninflamed (grey) and 2 inflamed tonsil donors (cyan). B) Proportion of cell phenotypes identified in tonsil from 3 inflamed (3 ROIs) and 6 noninflamed (6 ROIs) donors. C) Absolute count of NK cell subsets per tonsil domain from either 3 inflamed or 6 noninflamed donors used for CyCIF. D) Frequency of NK cell subsets from 9 inflamed and 6 noninflamed tonsil donors, significance was calculated by two-way ANOVA with multiple comparisons. E) Shortest median distance measurement of NK cell subsets to dendritic cells, CD68+ macrophages, and other myeloid cells quantified from CyCIF data of 3 inflamed and 6 noninflamed donors.

Supp. Figure 7

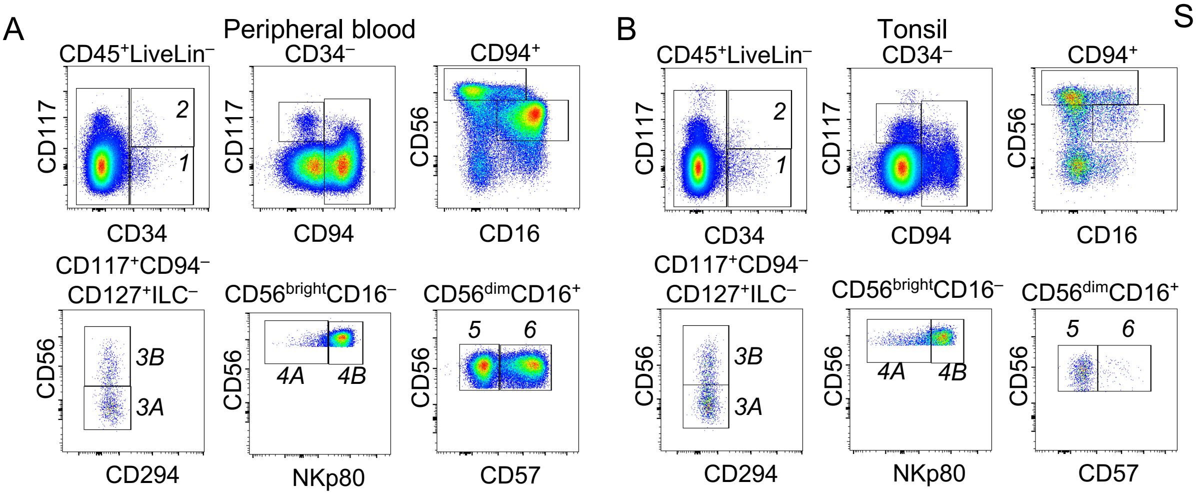

Supplemental Figure 7: A) Representative flow cytometry gating strategy for measuring peripheral blood and (B) tonsil NK cell subset chemokine receptor expression.

| Metal | Marker | Vendor | Catalog # | Clone/Identifier | Dilution |
|---|---|---|---|---|---|
| 115IN | GZMA | ABCAM | AB209205 | EPR20161 | 1:50 |
| 141Pr | SMA | Standard Biotools [Fluidigm] | 3141017D | 1A4 | 1:80 |
| 142Nd | GZMK | LSBio | LS-C119554 | Polyclonal | 1:50 |
| 143Nd | CD103 | ABCAM | AB224202 | EPR22590-27 | 1:50 |
| 144Nd | CD57 | Biolegend | 395602 | HNK1 | 1:40 |
| 145Nd | TBET | Biolegend | 644802 | 4B10 | 1:50 |
| 146Nd | CD16 | Standard Biotools [Fluidigm] | 91H004 | EPR16784 | 1:100 |
| 147Sm | CD200R | LSBio | LS-C378947 | Polyclonal | 1:50 |
| 148Nd | Pan keratin | Standard Biotools [Fluidigm] | 3148020D | C11 | 1:100 |
| 149Sm | CD15 | Standard Biotools [Fluidigm] | 3149026D | W6D3 | 1:25 |
| 150Nd | CD203c | ABCAM | AB150558 | Polyclonal | 1:40 |
| 151Eu | CD31 | Standard Biotools [Fluidigm] | 3151025D | EPR3094 | 1:100 |
| 152Sm | CD45 | Standard Biotools [Fluidigm] | 3152016D | CD45-2B11 | 1:50 |
| 153Eu | CD94 | Proteintech | 13332-1-AP | Polyclonal | 1:40 |
| 155Gd | HLA-C | Abcam | AB193432 | Polyclonal | 1:25 |
| 156Gd | CD122 | ThermoFisher | MA5-23816 | 27302 | 1:40 |
| 158Gd | NKp46 | Innate Pharma | n/a | 8E5B | 1:25 |
| 159Tb | CD56 | Millipore | AB5032 | Polyclonal | 1:25 |
| 160Gd | NKG2A | Miltenyi | 130-122-329 | REA110 | 1:25 |
| 161Dy | CD20 | Standard Biotools [Fluidigm] | 3161029D | H1 | 1:100 |
| 162Dy | CD8a | Standard Biotools [Fluidigm] | 3162034D | C8/144B | 1:100 |
| 163Dy | Eomes | Millipore | AB2283 | Polyclonal | 1:100 |
| 164Dy | CD117 | Agilent/DAKO | A4502 | Polyclonal | 1:20 |
| 165Ho | NKp80 | Miltenyi | 130-126-475 | REA845 | 1:20 |
| 166Er | CD68 | LSBio | LS-B17033 | C68/684 | 1:50 |
| 167Er | Granzyme B | ABCAM | AB208586 | EPR20129-217 | 1:100 |
| 168Er | CD127 | Standard Biotools [Fluidigm] | 3168026D | EPR2955(2) | 1:100 |
| 169Tm | CD117 | RnD Systems | AF1356 | Polyclonal | 1:20 |
| 170Er | CD3 | Standard Biotools [Fluidigm] | 3170019D | Polyclonal | 1:100 |
| 171Yb | RORgt | Millipore | MABF81 | 6F3.1 | 1:100 |
| 172Yb | KI67 | Standard Biotools [Fluidigm] | 3172024B | B56 | 1:30 |
| 173Yb | CD10 | ThermoFisher | PA5-47075 | Polyclonal | 1:25 |
| 174 Yb | HLA-E | ABCAM | AB2216 | MEM-E/02 | 1:25 |
| 175Lu | CD34 | Novus Bio | NBP2-34713 | QBEnd/10 | 1:40 |
| 176Yb | CD11c | ABCAM | AB52632 | EPI347Y | 1:100 |
| 194Pt | Vimentin | Cell Signaling Technologies | 5741 | D21H3 | 1:100 |
| 191/193Ir | Intercalator | Standard Biotools [Fluidigm] | 201192B | n/a | 1:400 |

Supplemental Table 1: Antibody panel used to stain FFPE tonsil tissue for imaging mass cytometry. Antibodies not sourced from Standard Biotools [Fluidigm] were conjugated in house using Standard Biotools [Fluidigm] Maxpar Multimetal Antibody Labeling Kit (Cat. # 201300).

Supplemental table 2

| Round | Marker | Manufacturer | Catalog # | Clone | Removed | Ab Concentration (mg/mL) |
|---|---|---|---|---|---|---|
| 1 | CD56 | RnD | AF2408 | Polyclonal | | 2.5 |
| | CD34 | Novus | NBP2-34713 | QBEnd/10 | | 4 |
| | IL2RB | LSBio | LS-C401487-60 | Polyclonal | | 1.5 |
| 2 | CD49a | RnD | AF5676 | Polyclonal | | 1 |
| | NKp46 | Thermofisher | PA5-79720 | Polyclonal | | 5 |
| | gP38 | Biolegend | 395002 | LpMab-21 | | 5.5 |
| 3 | ICAM1 | RnD Systems | BBA17 | Polyclonal | | 6 |
| | CD20 | BD | 555677 | H1 | | 2.5 |
| | CD127 | Invitrogen | PA5-122138 | Polyclonal | | 5 |
| 4 | CD10 | RnD Systems | AF1182 | Polyclonal | | 6 |
| | CD45 | Invitrogen | 14-9457-82 | CD45-2B11 | | 5 |
| | CD49d | LSBio | LS-B3837-0.05 | Polyclonal | | 7.5 |
| 5 | CD117 | RnD | AF1356 | Polyclonal | Yes | 3 |
| | Ki67 | Fluidigm | 3172024B | B56 | | 5 |
| | CD94 | LSBio | LS-B14034 | Polyclonal | | 10 |
| 6 | CD3 | ABCAM | AB11089 | CD3-12 | Yes | 10 |
| | CXCL12 | RnD Systems | MAB350-SP | 79018 | | 8 |
| | CCL21 | MyBioSource | MBS9411525 | Polyclonal | | 4.8 |
| 7 | CCL3 | RnD Systems | AF-270-NA | Polyclonal | Yes | 10 |
| | CD68 | LSBio | LS-B17033-100 | C68/684 | | 10 |
| | CD3 | ABCAM | AB5690 | Polyclonal | | 1.5 |
| 8 | CX3CL1 | RnD Systems | AF537 | Polyclonal | | 6 |
| | Flt3L | ABCAM | AB52648 | EP1140Y | | 4 |
| | NKp46 | RnD Systems | MAB1850 | 195314 | Yes | 8 |
| 9 | CCL4 | ThermoFisher | MA5-43912 | A4 | Yes | 7.5 |
| | IL7 | Sigma Atlas | HPA019590 | Polyclonal | | 2 |
| | VCAM1 | Novus | NBP2-44615 | VCAM1/843 | Yes | 3 |
| 10 | CD56 | RnD | AF2408 | Polyclonal | Yes | 3 |
| | CCL20 | ThermoFisher | MA5-43912 | A4 | | 10 |
| | GZMK | Invitrogen | PA5-84586 | Polyclonal | | 1.5 |
| 11 | CCL3 | RnD Systems | AF-270-NA | Polyclonal | Yes | 5 |
| | CD34 | Novus | NBP2-34713 | QBEnd/10 | Yes | 5 |
| | GZMB | ABCAM | AB4059 | Polyclonal | | 10 |
| 12 | CX3CL1 | RnD Systems | AF537 | Polyclonal | Yes | 4 |
| | CCL19 | ThermoFisher | TA804261S | OTI2A12 | Yes | 5 |
| | CD94 | ABCLONAL | A6438 | Polyclonal | | 10 |
| 13 | CD117 | RnD | AF1356 | Polyclonal | Yes | 2 |
| | CD122 | LSBio | LS-C401487-60 | Polyclonal | | 3 |
| 14 | ICAM1 | RnD Systems | BBA17 | Polyclonal | | 5 |
| | VCAM1 | Novus | NBP2-44615 | VCAM1/843 | Yes | 5 |
| | NCR3 | ABCLONAL | A14522 | Polyclonal | | 3.927 |
| 15 | CCL4 | ThermoFisher | MA5-43912 | A4 | Yes | 10 |
| | Pan Keratin | Fluidigm | 3148020D | C11 | | 5 |
| | IL15 | LSBio | LS-B10060 | Polyclonal | | 15 |
| 16 | SMA | Fluidigm | 3141017D | 1A4 | | 5 |
| | Collagen type 1 | Fluidigm | 3169023D | Polyclonal | | 3.75 |
| | CD69 | ABCLONAL | A21174 | Polyclonal | | 15 |
| 17 | CD57 | Biolegend | 359602 | HNK1 | | 15 |
| | CD117 | RnD | AF1356 | Polyclonal | | 4 |
| | CD103 | ABCLONAL | A9934 | Polyclonal | | 14.2 |
| 18 | Ki67 | Fluidigm | 3172024B | B56 | Yes | 6 |
| | CD56 | RnD | AF2408 | Polyclonal | Yes | 5 |
| | CD11c | ABCAM | AB52632 | EPI347Y | Yes | 7 |
| 19 | NKp46 | Innate Pharma | n/a | 8E5B | | 0 |
| | CD11c | Sigma Atlas | HPA004723 | Polyclonal | | 20 |
| | CX3CL1 | RnD Systems | AF537 | Polyclonal | Yes | |
| 20 | CCL19 | ThermoFisher | TA804261S | OTI2A12 | | 2 |
| | CD31 | Novus | NB100-2284 | Polyclonal | | 0.6 |
| | NKp80 | Miiltenyi | 130-112-591 | REA845 | | 20 |
| 21 | CD122 | LSBio | LS-C401487-60 | Polyclonal | Yes | 10 |
| 20 | TBET | Biolegend | 644802 | 4B10 | Yes | 10 |
| | CD127 | Invitrogen | PA5-122138 | Polyclonal | Yes | 7 |
| 23 | CD94 | Proteintech | 13332-1-AP | Polyclonal | | 15 |
| | CD8a | Fluidigm | 3162034D | C8/144B | | 5 |

Supplemental Table 2: Antibody panel used for immunophenotyping tonsil cell populations by CyCIF. 43 unique markers were used over 23 cycles of staining and imaging. 21 channels were removed for downstream analysis because they were either repeated stains or did not work based on low signal to noise ratio.

| Fluorophore | Marker | Clone/Identifier | Vendor | Catalog # |
|---|---|---|---|---|
| BUV395 | CD94 | HP-3D9 | BD Biosciences | 743954 |
| BUV496 | CD3 | UCHT1 | BD Biosciences | 612940 |
| BUV496 | CD14 | M5E2 | BD Biosciences | 750381 |
| BUV496 | CD19 | SJ25C1 | BD Biosciences | 612938 |
| BUV563 | CD56 | NCAM16.2 | BD Biosciences | 612928 |
| BUV805 | CD45 | HI30 | BD Biosciences | 612891 |
| BV510 | CD294 | BM16 | Biolegend | 350120 |
| BV650 | NKp44 | p44-8 | BD Biosciences | 744302 |
| BV711 | CD117 | 104D2 | Biolegend | 313230 |
| BV785 | CD103 | Ber-ACT8 | Biolegend | 350230 |
| PE | Ki67 | KI67 | Biolegend | 350504 |
| PE/Dazzle™ 594 | CD34 | 561 | Biolegend | 343534 |
| PE-Cy5 | CD127 | A019D5 | Biolegend | 351324 |
| APC | NKp80 | REA845 | Miltenyi | 130-112-591 |
| AF700 | CD16 | 3G8 | Biolegend | 302026 |
| APC-Cy7 | Zombie NIR | n/a | Biolegend | 423106 |

Supplemental Table 3: Antibody panel used to immunophenotype proliferating NK cell subsets from tonsil by flow cytometry.

| Panel | Fluorophore | Marker | Clone/Identifier | Vendor | Catalog # |
|---|---|---|---|---|---|
| all | BUV395 | CD94 | HP-3D9 | BD Biosciences | 743954 |
| all | BUV496 | CD3 | UCHT1 | BD Biosciences | 612940 |
| all | BUV496 | CD14 | M5E2 | BD Biosciences | 750381 |
| all | BUV496 | CD19 | SJ25C1 | BD Biosciences | 612938 |
| all | BUV563 | CD56 | NCAM16.2 | BD Biosciences | 612928 |
| 1 | BV421 | CXCR3 | G025H7 | Biolegend | 353715 |
| 2 | BV421 | CCR5 | J418F1 | Biolegend | 359117 |
| 3 | BV421 | CCR7 | G043H7 | Biolegend | 353208 |
| 4 | BV421 | CXCR4 | 12G5 | Biolegend | 306517 |
| all | BUV805 | CD45 | HI30 | BD Biosciences | 612891 |
| all | BV510 | CD294 | BM16 | Biolegend | 350120 |
| all | BV650 | NKp44 | p44-8 | BD Biosciences | 744302 |
| all | BV711 | CD117 | 104D2 | Biolegend | 313230 |
| all | BV785 | CD103 | Ber-ACT8 | Biolegend | 350230 |
| all | FITC | CD57 | HNK-1 | Biolegend | 359604 |
| 1 | PE | CCR4 | G034E3 | Biolegend | 353418 |
| 2 | PE | CCR9 | L053E8 | Biolegend | 358903 |
| 3 | PE | CCR8 | L263G8 | Biolegend | 360603 |
| 4 | PE | CXCR5 | J252D4 | Biolegend | 356904 |
| all | PE/Dazzle™ 594 | CD34 | 561 | Biolegend | 343534 |
| all | PE-Cy5 | CD127 | A019D5 | Biolegend | 351324 |
| 1 | PE-Cy7 | CCR6 | G034E3 | Biolegend | 353418 |
| 2 | PE-Cy7 | CCR1 | 5F10B29 | Biolegend | 362913 |
| 3 | PE-Cy7 | CX3CR1 | 2A9-1 | Biolegend | 341611 |
| 4 | PE-Cy7 | CXCR6 | K041E5 | Biolegend | 356011 |
| all | APC | NKp80 | REA845 | Miltenyi | 130-112-591 |
| all | AF700 | CD16 | 3G8 | Biolegend | 302026 |
| all | APC-Cy7 | Zombie NIR | n/a | Biolegend | 423106 |

Supplemental Table 4: Antibody panels used to measure NK cell expression intensity of chemokine receptors by flow cytometry. 4 panels were used to measure 12 chemokine receptors; all panels included a single backbone of NK cell markers used to identify all subsets. 6-7 paired tonsil and peripheral blood donors were used per panel.

| Fluorophore | Marker | Clone/Identifier | Vendor | Catalog # |
|---|---|---|---|---|
| BUV496 | CD3 | UCHT1 | BD Biosciences | 612940 |
| BUV496 | CD14 | M5E2 | BD Biosciences | 750381 |
| BUV496 | CD19 | SJ25C1 | BD Biosciences | 612938 |
| BUV563 | CD56 | NCAM16.2 | BD Biosciences | 612928 |
| BUV805 | CD45 | HI30 | BD Biosciences | 612891 |
| Pacific Blue | CD31 | WM59 | Biolegend | 303124 |
| BV650 | gp38 (podoplanin) | LpMab-17 | BD Biosciences | 747633 |
| BV711 | CD73 | AD2 | Biolegend | 344026 |
| FITC | CD271 | ME20.4 | Biolegend | 345104 |
| PE | CD56 | 39D5 | Biolegend | 355504 |
| PE/Dazzle™ 594 | CD34 | 561 | Biolegend | 343534 |
| PE-Cy5 | CD106 | STA | Biolegend | 305808 |
| PE-Cy7 | CD90 | 5-E10 | Biolegend | 328124 |
| APC | CD10 | HI10A | Biolegend | 312210 |
| AF700 | CD54 | HA58 | Invitrogen | 56-0549-42 |
| APC-Cy7 | ZombieNIR | n/a | Biolegend | 423106 |

Supplemental Table 5: Antibody panel utilized for immunophenotyping bone marrow and tonsil stromal cell populations.

| Sample ID | Sex | Age (years) | Inflamation | Pathology Report | Patient notes |
|---|---|---|---|---|---|
| ENT057 | M | 4.75 | yes | acute tonsilitis and lymphoid hyperplasia | |
| ENT059 | M | 4.67 | yes | acute tonsilitis and lymphoid hyperplasia | |
| ENT061 | F | 3.67 | no | lymphoid hyperplasia | |
| ENT066 | M | 3.42 | yes | chronic and acute tonsillitis | |
| ENT067 | M | 4.25 | no | | |
| ENT069 | M | 6.67 | no | | |
| ENT079 | M | 4.9 | no | | |
| ENT088 | M | 6.9 | no | | |
| ENT089 | M | 7.5 | no | | |
| ENT091 | F | 4.5 | yes | | recurring throat infections |
| ENT100 | F | 17 | no | Hypertrophy of tonsils and adenoids | obstructive sleep apnea |
| ENT128 | M | 8.8 | no | Hypertrophy of tonsils and adenoids | recurring throat infections |

Supplemental Table 6: Pediatric donors used for CyCIF and IMC analysis.
